# Back-projection improves inference from sparsely sampled genomic surveillance data

**DOI:** 10.1101/2025.06.29.662219

**Authors:** Elizabeth E. Finney, Brian Lee, Syed Faraz Ahmed, Muhammad Saqib Sohail, Ahmed Abdul Quadeer, Matthew R. McKay, John P. Barton

## Abstract

Highly transmissible SARS-CoV-2 variants have emerged throughout the COVID-19 pandemic, driving new waves of infections. Genomic surveillance data can provide insights into the virus’s evolution and biology. However, delayed and limited regional data can introduce biases in epidemiological models, potentially obscuring transmission patterns. To address this issue, we used a novel, variant-specific back-projection model to estimate a distribution of likely infection times from sample collection times. We combined this approach with epidemiological modeling to estimate selection for increased transmission in a way that accounts for the uncertainty in infection times. Tests in simulations demonstrated that our method can make the inference of selection more reliable. We also applied our approach to SARS-CoV-2 data, where it excelled in smoothing and extending data from geographic regions or times with poor sampling. Overall, our method can aid in the reliable identification of mutations and variants with higher transmission rates.

## Introduction

The emergence of new viral variants can pose a major challenge to public health. New variants can be more transmissible than past ones or have antigenic changes that allow them to escape natural or vaccine-induced immunity. Multiple such variants of concern (VOCs) and variants of interest (VOIs) have arisen throughout the COVID-19 pandemic^1,2^. The spread of new viral variants is also relevant for other viruses such as influenza, where the accumulation of mutations in surface proteins enables the virus to escape control by antibodies^3,4^. Detecting and characterizing these variants could improve outbreak responses and reveal key mechanisms of viral adaptation.

Genomic surveillance data provides insights into viral evolution, but several factors complicate the analysis of variant transmission. Data can contain regional and spatial limitations, time delays that occur between infection and sequence collection, and gaps in coverage due to under-reporting or other effects. These all introduce biases when modeling viral spread^5–11^. In particular, reported sequence collection times may be delayed compared to actual infection times, obscuring underlying patterns of transmission.

To address the challenges of incomplete and delayed surveillance data, epidemiologists have developed statistical methods to reconstruct patterns of disease transmission. One of these methods is back-projection, an approach that models the underlying rates of disease transmission over time based on noisy, incompletely observed samples. This method was first developed during the HIV/AIDS epidemic, where long delays between infection and the development of AIDS made it difficult to track the spread of disease in real time^12^. Based on an expectation maximization-smoothing (EMS) approach, the original back-projection algorithm proved highly effective at reconstructing HIV infection dynamics from AIDS case data. This method has since been adapted to study outbreaks of other diseases, including the original outbreak of SARS in 2003, anthrax infections, and the early spread of SARS-CoV-2^10,13–17^.

Here, we applied back-projection methods to improve studies of viral evolution. As a novel feature of our approach compared to prior work, we adapted back-projection to infer the transmission dynamics of individual variants rather than treating all cases as equivalent. We then combined back-projected estimates of variant frequencies with evolutionary modeling^18,19^ to estimate how different mutations affect viral transmission. Tests in simulations show that our approach makes estimates of mutational effects from finitely sampled data more robust. We then applied our method to study SARS-CoV-2 evolution using genomic surveillance data from GISAID. In geographical regions with infrequent sampling, our method allows us to reconstruct variant frequencies and transmission patterns that would otherwise be obscured by gaps in surveillance. This ability to extract reliable information from limited data is particularly valuable as SARS-CoV-2 clinical sequencing efforts continue to decline^20,21^, and as researchers turn to alternate surveillance strategies to aid clinical testing^22,23^. Amid changing surveillance strategies, this approach could help maintain effective genomic surveillance with fewer resources.

## Results

### Estimating infection rates with back-projection

First, we will introduce the back-projection method^12,29^. For each viral variant *a*, we write the true number of infections that occur during some discrete interval of time *i* as *N*_*i*_(*a*). The time interval should be chosen appropriately for the problem of interest; for SARS-CoV-2, we will measure time in days. However, the true number of infections is not directly observed. Instead, there are a certain number of recorded viral sequences *Y*_*i*_(*a*) during each time interval *i*. The true number of infections and recorded sequences are then connected by a random process, depending on the time between infection and the development of symptoms and testing for infection, and the probability that the virus that an infected individual harbors is sequenced.

The time between infection and the development of symptoms is called the incubation period (**Fig. 1**). Incubation periods are stochastic and are thus described by a distribution, rather than a single time. We quantify the incubation period distribution by variables *f*_*i*_ that give the probability that the incubation period is *i* days. For simplicity, we will merge the probabilities of testing and sequencing into the incubation period distribution, such that this gives the probability that the virus infecting an individual will be sequenced *d* days after infection.

**Fig. 1.**
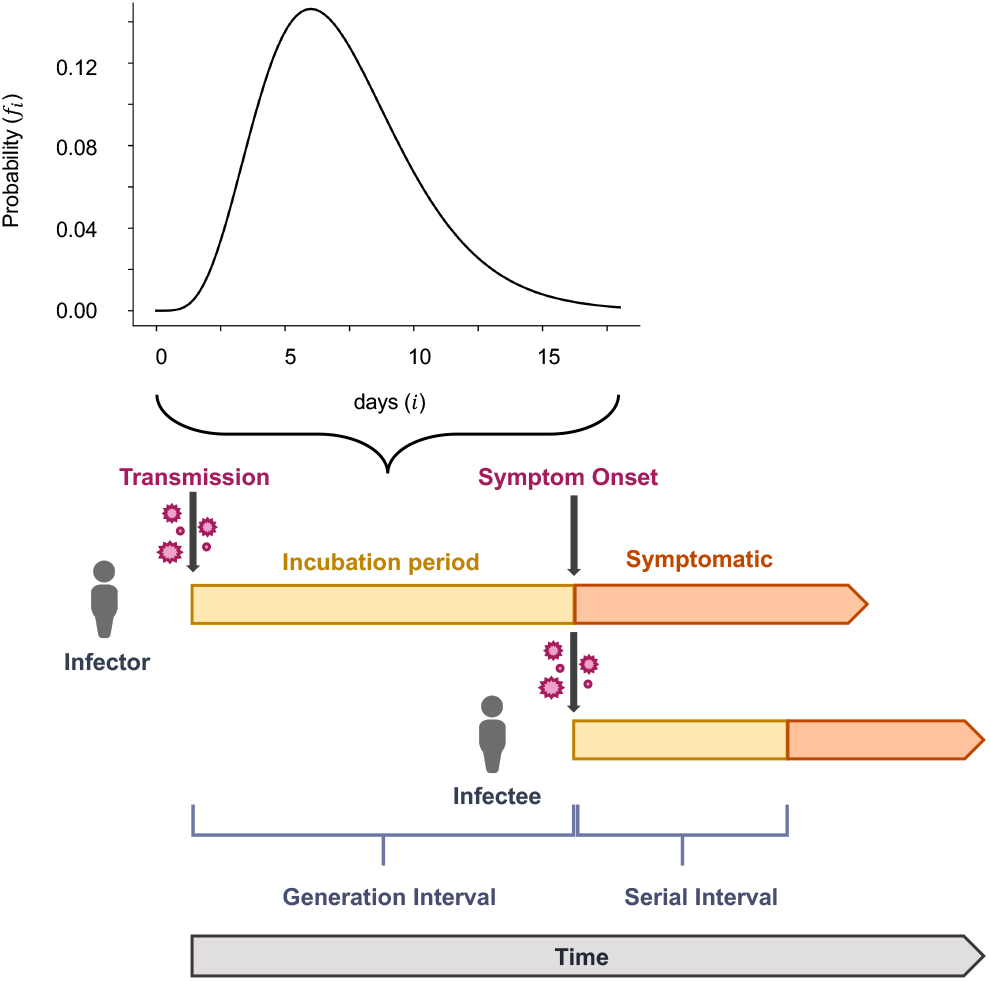
Illustration of transmission. When an individual (the infector) is infected with a virus, an incubation stage occurs before they are able to spread the virus to another individual (the infectee). We assume that samples are likely to be collected near the onset of symptoms, which is also close to the peak of transmissibility for SARS-CoV-2^24,25^. This is equivalent to using the incubation period as a proxy for the generation interval (the time interval between infection onset in an infection pairing). We make this assumption because the generation interval is rarely observable^26^ and is often not well approximated by using the serial interval (the time interval between symptom onset in an infection pairing) as a proxy instead^27^. The SARS-CoV-2 incubation period has been modeled by a Gamma distribution, where the median incubation period is around 5 days^28^.

Of course, most individuals who are infected with SARS-CoV-2 or other pathogens do not undergo sequencing. Thus, there is also a high probability that the virus that infects an individual is never sampled. We are concerned with the relative prevalence of different viral variants, so we will not model the probability that an infected individual is not sequenced. While this probability is highly relevant for estimating underlying numbers of infections, it does not affect the estimated proportion of infections due to one viral variant or another.

Now, we wish to estimate the true rate of infections 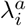 (proportional to *N*_*i*_(*a*)) for variant *a* at each time *i* from the observed infections ***Y*** (*a*) = (*Y*_1_(*a*), *Y*_2_(*a*),…, *Y*_*T*_ (*a*)) and the incubation time distribution. Becker, Watson, and Carlin provided an approximate solution to this problem using an expectation maximization approach^12^, which we describe below. To simplify this problem, let us assume that the number of infections during some time *i* and with some incubation period *j*, which we write 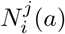, follows a Poisson distribution with rate 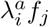. The likelihood of the 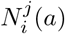, counting over all incubation times *j*, would then be given by

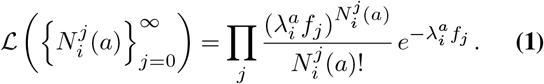

For each *i*, the 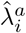 that maximizes the log-likelihood would then be found by solving

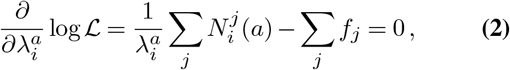

which yields

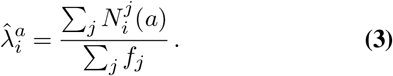

However, the 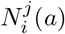 are unknown. Our data contains only the observed number of infections at each time ***Y*** (*a*). Let us assume that we knew the underlying incidence rates together with the data. Following the Poisson assumption above, we have 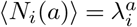. We can also connect our data with the 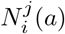, as 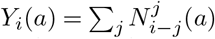. In general, for independent Poisson random variables *X*_*i*_ with rates *r*_*i*_ that are constrained to sum to *X*, the expected value of *X*_*i*_ is given by

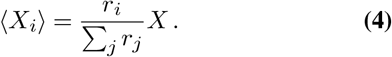

We can therefore write the expected value for the 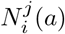, depending on the observations ***Y***(*a*) and the underlying infection intensities ***λ***^*a*^ as

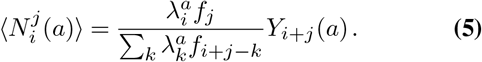

Thus, provided an initial estimate for the ***λ***^*a*^, (3) and (5) can be combined to provide an iterative formula for estimating the underlying infection intensities,

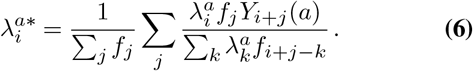

One then replaces the ***λ***^*a*^ with the ***λ***^*a∗*^ from (6), repeating this process until convergence (Methods). In practice, a smoothing step is also added to constrain 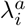 values from around the same time period to be similar to one another, encoding an implicit assumption that underlying infection rates do not change too rapidly in time (Methods). Compared to prior work, our estimation of infection intensities for individual variants allows us to reconstruct variant dynamics from finitely sampled data.

While reconstructing underlying infection intensities is, on its own, an interesting problem, we are also especially interested in using reconstructed infection intensities in down-stream analyses. Below, we describe one application: using smoothed infection intensities, rather than raw sample collection times, to infer how different mutations affect the transmission of a pathogen.

### Inferring the transmission effects of mutations

One key interest in studying the dynamics of viral variants is to identify new variants and mutations that alter transmissibility. Here, we applied a recently developed method to estimate the effects of individual mutations on viral transmission from genomic surveillance data^19,30^. Related methods have been applied to study evolution in a variety of different contexts, including the evolution of viruses within hosts^18,31–33^, bacteria^34,35^, and high-throughput experiments^36^. This method uses a branching process model of disease transmission where the effective reproduction number *R*_*a*_ depends on the viral variant *a* that an individual is infected with. Specifically, Lee et al. assume a simple, additive model for the effects of viral mutations on transmission, such that

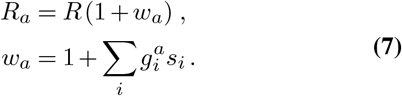

Here, the *w*_*a*_ and *s*_*i*_ are referred to as selection coefficients in analogy with models in evolutionary biology. The *w*_*a*_ quantifies how much more or less transmissible viral variant *a* is compared to the reference, which is chosen to have an effective reproduction number of *R* by convention. Differences in transmissibility are ultimately explained by the effects of individual mutations *s*_*i*_, where *i* labels a specific mutation. The 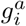 are indicator variables which are equal to one if variant *a* has the mutation *i* and zero otherwise.

To estimate the selection coefficients from genomic surveillance data, Lee et al. compute the probability of different evolutionary trajectories of the virus population as a function of the selection coefficients. One can then derive the selection coefficients that best explain the evolutionary history of the virus, in the sense that they would have the highest probability to produce the observed data^19^ (Methods). To combine this approach with back-projection, we replace the noisy empirical variant frequencies with our estimated infection intensities ***λ***^*a*^. Intuitively, variants or mutations that increase transmission are likely to represent a larger fraction of infections over time, while those that are less competitive are likely to decline in frequency. The more rapidly a variant or mutation increases or decreases in frequency, the larger the estimated change in transmission will typically be.

### Validation in simulations

We first tested our back-projection and inference approach in simulations. We simulated the spread of six different viral variants among a population of susceptible hosts. Two of the variants had beneficial mutations (*w* = 3%), two had neutral mutations (*w* = 0), and two had deleterious mutations (*w* = −3%). We assumed stochastic transmission dynamics where the number of new infections generated by one infected individual follows a negative binomial distribution. The negative binomial distribution is heavy-tailed and has been used in past work to account for superspreading^37–40^. Here, we set the dispersion parameter *k* = 0.1 and adjusted the overall effective reproduction number *R* such that the number of infected individuals in the simulation was roughly constant over time (Methods). To thoroughly test the typical results for back-projection, we performed 10^4^ simulations with the same underlying parameters.

To mimic variation in the development of symptoms and sample collection, we drew the observed sample times (measured after the true transmission time) from a gamma distribution with shape and scale parameters of 5.8 and 0.95, respectively. These parameters match with early estimates of the incubation period of SARS-CoV-2 (ref.^28^). In this way, the sample times in our simulations are delayed, “blurred” versions of the true transmission times.

We then applied back-projection to the simulated data to reconstruct the underlying transmission processes (**Fig. 2**). As noted above, we estimated separate infection intensities for each variant rather than grouping all variants together. We used binomial smoothing with a window size of 8 days, limiting the maximum number of iterations to 250 and setting a stopping criterion of *ϵ* = 10^*−*4^ for the convergence of the infection intensities ***λ*** (refs.^12,14,41^). We also truncated the ends of the trajectories by eliminating time points where the summed intensities of all variants were less than 20, similar to procedures used to exclude periods of poor sampling in naive analyses without smoothing^19^. As shown in **Fig. 2**, back-projection produces smoothed versions of the true infection histories.

**Fig. 2.**
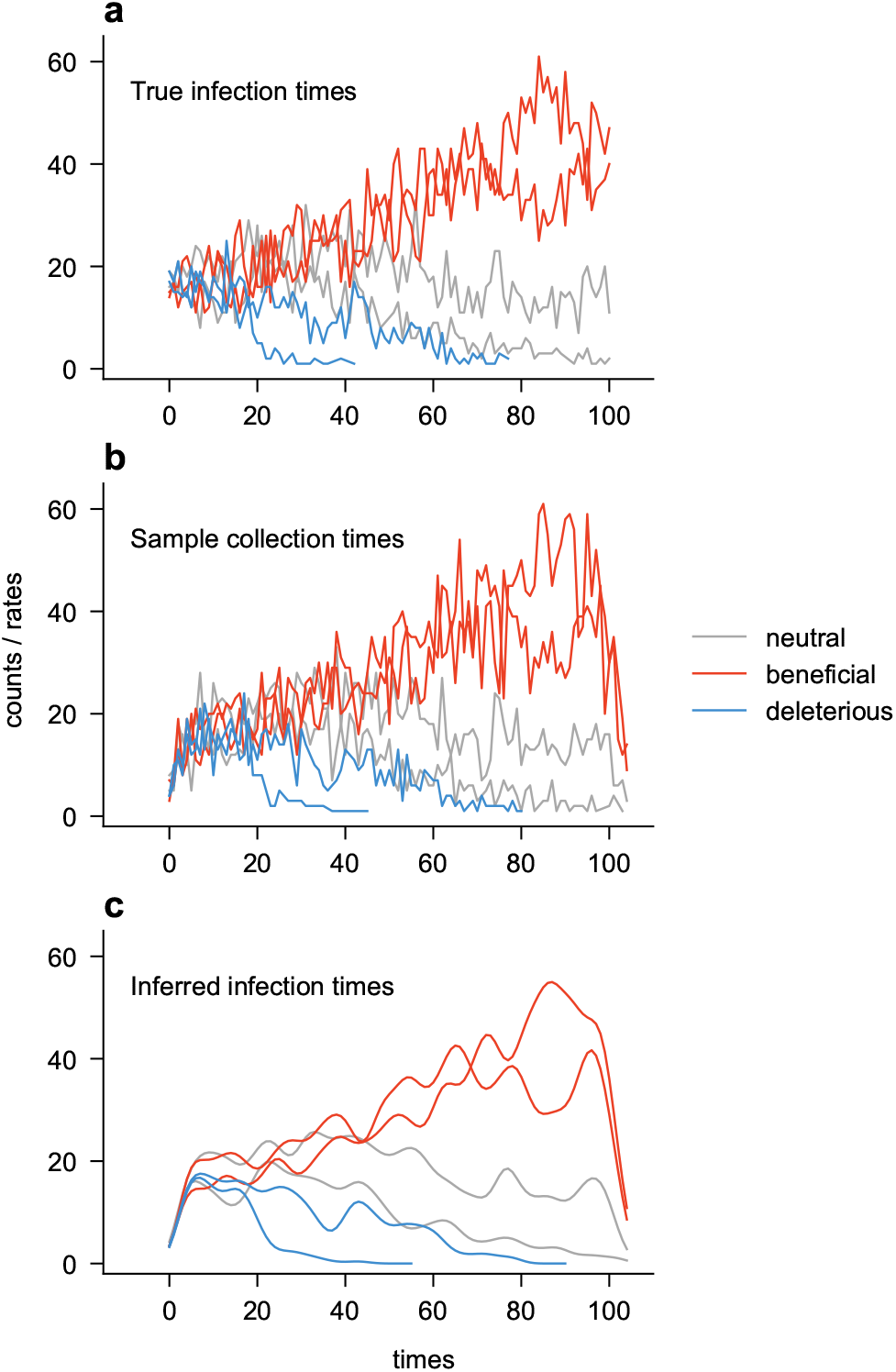
Simulated infection histories. Example simulation of the spread of infection for six variants according to a branching process (Methods), with two each having neutral (*w* = 0), beneficial (*w* = 3%), and deleterious (*w* = *−*3%) mutations. **a**. True infection histories. **b**. Sampled infection histories, assuming that the shifts in the sampling time relative to the infection time follow a Gamma distribution. **c**. Smoothed infection histories recovered by back-projection from the sampled ones.

To test the utility of back-projection for improving our understanding of differences in transmission between variants, we inferred selection coefficients for each variant in the simulations. We performed inference using three different types of data. First, we inferred selection coefficients using the true transmission times in the simulations, which should yield the most accurate estimates. We also inferred selection coefficients using the simulated sample collection times and the back-projected infection intensities. In principle, inference using sample collection times should be less accurate than inference using the back-projected intensities because of the blurring of the infection times, which back-projection attempts to correct.

The simulations that we considered are challenging ones for demonstrating the effect of back-projection on downstream analyses. Here, we assumed that sequence data was abundantly sampled from the population. While this provides large amounts of data to infer infection intensities, the original data is already so well-sampled that smoothed estimates of underlying infection rates may be unnecessary to infer how different variants affect transmission. In our applications to real SARS-CoV-2 data below, we will also explore scenarios where sampling is much sparser. These cases show more dramatic differences between data analysis with and without back-projection, including significant variation in the length of the sample trajectories, which is not present in our well-sampled simulations.

**Figure 3** shows that, even in this well-sampled case, back-projection can improve the inference of transmission effects from surveillance data compared to analysis performed using sample collection times. Inferences using back-projection are similar to those using the true infection times. Back-projection is especially helpful for controlling occasional large errors in inferences using raw sample collection times (see tails of the distribution in **Figs. 3a, c**). Compared to inference using the true infection times, back-projection actually offered slight improvements in the ability to reliably classify variants as beneficial or deleterious (i.e., increasing or decreasing transmission, respectively), which may be attributed to smoothing of the infection intensity parameters.

**Fig. 3.**
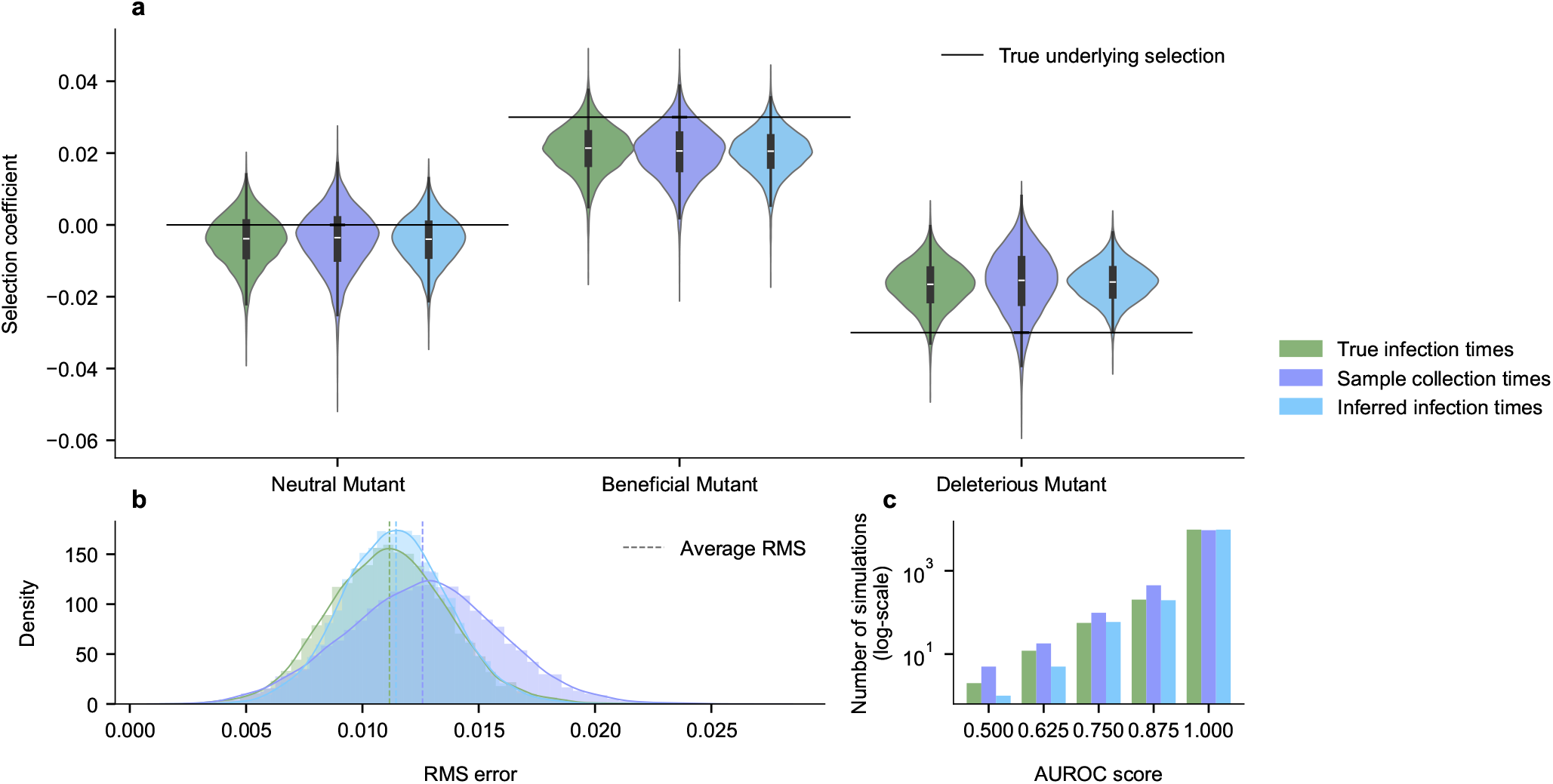
Inferred transmission effects of mutations in simulations. **a**. Distribution of inferred selection coefficients across 10^4^ simulations. Compared to using the empirically sampled frequencies, selection coefficients inferred with back-projected infection intensities take on less extreme values and are closer to the true, underlying ones. **b**. Distributions of root mean squared (RMS) errors of the inferred selection coefficients show that back-projection reduces average errors compared to inferences with raw sample collection times. **c**. Comparison of area under the receiver operating characteristic (AUROC) scores that measure the accuracy of classifying mutations that increase viral transmission from data. Higher values indicate more successful classification. With our simulation parameters, misclassification is rare (*∼* 5% of simulations). By this metric, inference from back-projected data even slightly outperforms the true infection frequencies (as there are fewer cases with misclassifications), which may be attributable to smoothing.

### Back-projection for SARS-CoV-2 data

We applied back-projection to estimate SARS-CoV-2 infection histories from sampled sequence data from the GISAID repository^43^. Sequence data used in our analysis were collected between January 1, 2020, and January 26, 2024. Following prior data processing conventions^19^, we sorted sequence data into medium-sized geographical regions. These mostly corresponded to states or provinces in large countries such as the United States of America and entire countries in areas such as Europe. Since we expect back-projection to be most useful when data is limited, we focused our analysis on 30 regions with fairly sparse sampling (Methods). As we had done in simulations, we inferred separate infection intensities for each unique viral variant in the data, across each geographical region (**Supplementary Fig. 1**).

To perform back-projection, we used the same Gamma-distributed incubation time distribution described above, with shape and scale parameters of 5.8 and 0.95 (ref.^28^). These parameters were estimated from early SARS-CoV-2 transmission data. Other estimates of the incubation time distribution have also been performed, and there is some evidence that it may be different for different variants^44–46^. In principle, delays in sample collection could also be different in different regions. Here, for simplicity we keep the incubation time distribution parameters fixed across different regions and times.

Compared to prior studies without data smoothing^19^, back-projection provided smoothed estimates of underlying infection intensities that improved the resolution of data for regions and times with sparse sampling. To avoid spurious inferences, Lee et al. excluded data from times when *<* 20 sequences were collected over a window of 5 days. Applying the same criterion to infection intensities rather than raw counts, we found that we could extend the range of time that we analyze for most regions, by an average of around 19 days (**Supplementary Fig. 2**). **Figure 4** shows a typical example where the beginning of the sample range is extended. As a result, the rise of the B.1.1.63 variant in Hong Kong during their third wave of infection^47^ is recovered. The application of back-projection to data from South Africa is especially helpful for uncovering hidden evolutionary dynamics, as this adds around 3 weeks of data covering the rise of Omicron in late 2021 that would otherwise be excluded by strict sampling criteria (**Fig. 5**).

**Fig. 4.**
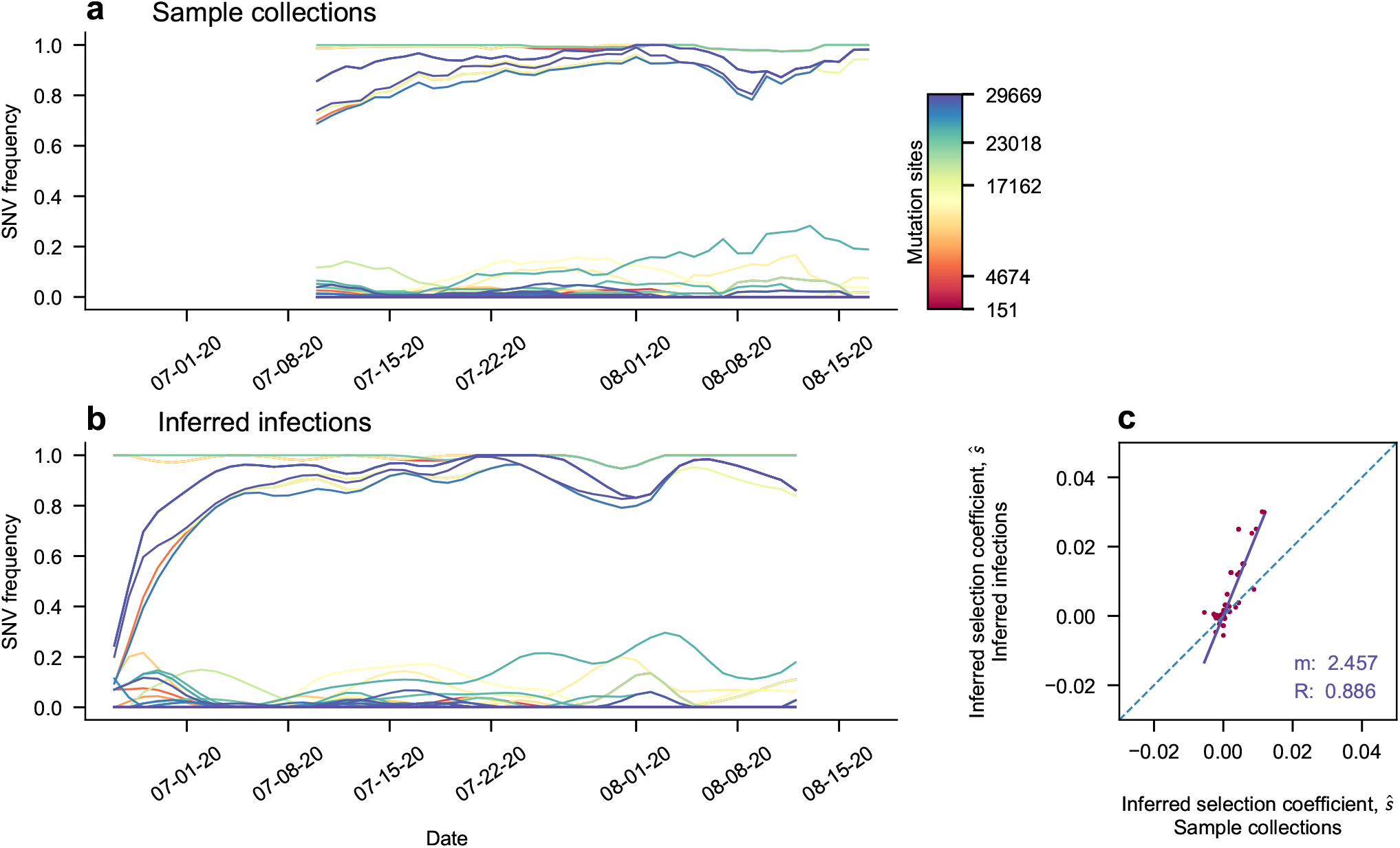
Evolutionary Trajectories During an Outbreak in Hong Kong, and Selection Coefficient Comparison. An example of a typical result of smoothing frequency trajectories via back projection. **a**, Unsmoothed trajectories of reported transmissions. **b**, Trajectories of inferred infections display smooth curves with variation corresponding with a typical shape of a propagating outbreak^42^, governed by the serial interval of the virus. There is newly visible left side behavior. **c**, The evolutionary histories affect the spread of selection coefficients inferred. By plotting unprocessed inferred selection coefficients against those inferred after back projection was applied, coefficients are pushed to more extreme values. They are inferred to be more beneficial if they were beneficial before processing, or more deleterious if they were deleterious. The process of smoothing and added evolutionary context could add confidence to these selection estimates. The purple line shows the linear regression fit of the selection coefficient comparisons, with a slope of 2.457, and a Pearson correlation coefficient of 0.886. The 1 to 1 line is shown as a dotted blue, for reference.

**Fig. 5.**
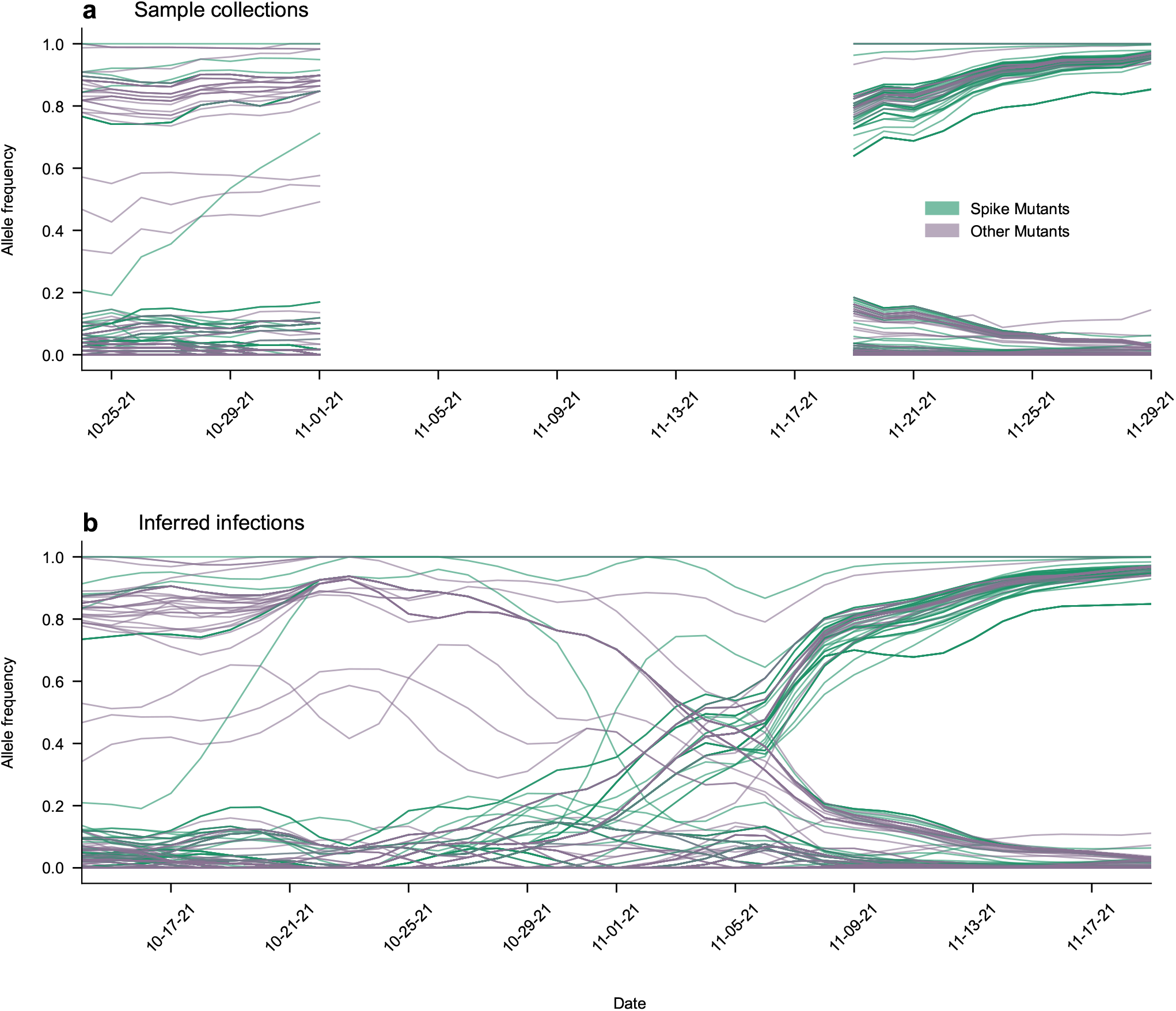
Back Projection Recovers Missing Information: Trajectories of SNVs during the rise of Omicron in South Africa. A period of poorly sampled sequences in the South African region happens to coincide with the rise of the Omicron variant. Due to these unreliable data, we were previously unable to examine the histories of mutations undergoing this selective sweep. Applying back projection allows for greater recovery of this information, leading to better estimates of evolution.

Next, we considered how back-projection affects the estimated effects of SARS-CoV-2 mutations on viral transmission^19^. Back-projection can affect these estimates by both increasing the amount of data available for analysis and by substituting smoothed infection intensities for more noisy empirical frequencies. **Figure 4c** shows a typical comparison between the inferred effects of SARS-CoV-2 mutations on viral transmission using back-projected and raw data. The selection coefficients shown here are inferred from the evolutionary trajectories shown in **Fig. 4a,b**. The two sets of selection coefficients are significantly correlated, but those inferred using back-projection are often somewhat larger in magnitude due to the larger time range of data included in the analysis. This provides greater evidence to support large positive or negative mutational effects on transmission. **Supplementary Fig. 2** shows that selection coefficients inferred with and without back-projection are typically highly correlated when the lengths of the trajectories with and without back-projection are similar (e.g., *R* ∼0.86 when trajectory lengths differ by 20% or less; additional examples are given in **Supplementary Figs. 3-18**).

## Discussion

In this work, we combined back-projection with evolutionary modeling to better understand the transmission dynamics of viral variants. Our approach differs from traditional back-projection by estimating infection histories for individual variants rather than grouping all variants together. Tests in simulations demonstrated that back-projection improves our ability to estimate mutational effects on viral transmission from finitely sampled, noisy data. When applied to SARS-CoV-2 surveillance data, our method proved particularly valuable for analyzing regions with sparse sampling, where it allowed us to reconstruct variant frequencies and transmission patterns that would otherwise be obscured by relative gaps in surveillance.

Our approach provides a principled way to handle sparse sampling and delays in genomic surveillance data. The expectation maximization algorithm used to solve the back-projection problem is mathematically rigorous and well-suited to handling incomplete data. In our case, “incompleteness” includes samples of variants that are delayed and uncertain relative to their true infection times. Adding a smoothing step makes the method more robust to outliers and missing data, which are common in real surveillance efforts. These features allow us to reconstruct transmission dynamics even in geographical regions or time periods where sampling is relatively sparse.

However, our model also has important limitations. Without additional information, we cannot explicitly correct for region-specific variations in sampling delays or changes in testing practices over time. While these issues helped to motivate our use of a robust smoothing method, a more detailed solution would require careful investigation of testing protocols in specific regions. Our focus on medium-sized geographical regions and population-level dynamics also means that we ignore fine-scale transmission patterns that might be captured by network or agent-based models^48,49^. Like any method that relies on surveillance and publicly available data sets, we are limited by the scope and frequency of data collection, which do not provide a complete picture of infection dynamics^50–53^.

The method also faces computational challenges when analyzing regions with extremely dense sampling. Back-projection effectively redistributes each discrete sequence across multiple potential infection times according to the incubation period distribution. This “blurring” of sequences across time points increases the computational burden of estimating the transmission effects of mutations compared to a naive approach where sample collection times are treated as exact. As a result, while our approach is well-suited for regions with sparse or moderate sampling density, it becomes computationally intensive for regions with very large numbers of sequences, such as the United Kingdom for SARS-CoV-2. However, improved methods for estimating underlying infection rates are also not as important for regions with extensive surveillance programs.

The COVID-19 pandemic has seen dramatic changes in testing and surveillance practices over time^11,54^. In general, SARS-CoV-2 testing has declined substantially from its peak, though test positivity rates remain high^20^. This decline stems from multiple factors: a shift from laboratory-based PCR testing to widespread use of rapid antigen tests^55,56^, changes in public health attitudes and behaviors^57–59^, and the impact of widespread vaccination^60,61^. While these developments represent important progress in managing COVID-19, they have also reduced the availability of viral sequences for tracking the emergence of new variants.

The ability to extract reliable information from limited data is especially valuable in regions or time periods where sequencing is infrequent. For example, applying our method to data from South Africa revealed transmission patterns during the emergence of the Omicron (BA.1) variant that would have been obscured under stringent data filtering criteria. As global SARS-CoV-2 sequencing efforts decline, methods that maximize the utility of available data will become increasingly important for maintaining effective genomic surveillance with fewer resources.

Future work could expand our method in several directions. The approach could be modified to explicitly account for regional differences in testing protocols and delays, though this would require a more detailed investigation of local surveillance practices. Our framework could also be adapted to analyze data from wastewater surveillance, which is becoming an increasingly important tool for monitoring viral spread at the community level^62^. Computationally, it may be advantageous to replace the ad hoc smoothing procedure used here and in prior work with a soft smoothness constraint on infection intensities. This change could allow for sharper changes in the underlying infection intensities when they are well-supported by data.

While we focused on SARS-CoV-2 evolution, our method is general and could be applied to study the transmission of other pathogens. The same framework could be used to analyze the spread of influenza variants or other viruses where surveillance data may be incomplete or delayed. Indeed, no virus has been sequenced as frequently or abundantly as SARS-CoV-2; gaps in pathogen surveillance data are ubiquitous. The key requirements of our approach are that the pathogen’s incubation period distribution is well-characterized and that some genomic surveillance data is available, even if sampling is sparse.

We have presented a method that draws from evolutionary biology, statistical physics, and epidemiological surveillance to better understand viral transmission. By making more effective use of limited surveillance data, our approach can help maintain insight into the spread of viral variants even as traditional sequencing efforts decline. This capability will be increasingly valuable for anticipating and controlling emerging viral threats.

## ACKNOWLEDGEMENTS

We gratefully acknowledge all data contributors, i.e., the Authors and their Originating laboratories responsible for obtaining the specimens, and their Submitting laboratories for generating the genetic sequence and metadata and sharing via the GISAID Initiative, on which this research is based. The work of E.F., B.L., and J.P.B. reported in this publication was supported by the National Institute of General Medical Sciences of the National Institutes of Health under Award Number R35GM138233. The work of A.A.Q., S.F.A. and M.R.M. was supported by the Australian Research Council (ARC) under Discovery Project, DP230102850. A.A.Q. and M.R.M. were also supported by the Research Grants Council of the Hong Kong Special Administrative Region, China under Project No. T11-705/21-N. M.R.M. is the recipient of an ARC Future Fellowship (Project No. FT200100928). The first author would like to acknowledge the new wave band, Talking Heads, for all of their musical support.

## AUTHOR CONTRIBUTIONS

All authors contributed to methods development, data analysis, interpretation of results, and writing the paper. E.F. led simulation and application of the back-projection approach to SARS-CoV-2 data. J.P.B. supervised the project.

## Methods

### Back-projection as a deconvolution method

The back-projection method can be thought of as a deconvolution applied to available case observations, in order to create a reconstruction of the infection process. We assume that infections occur according to a Poisson process with intensity *λ*_*t*_ at time *t*. Here for simplicity we drop the variant index *a*, which was shown explicitly in the main text. The fact that infection occurs via person to person transmission is ignored in this method, and we instead allow infections to occur by a random process so that *λ*_*t*_ is flexible enough to reproduce the infection intensity of a given transmission model^63,64^. Let *N*_*t*_ denote the number of individuals infected from time 0 to time *t*, so that

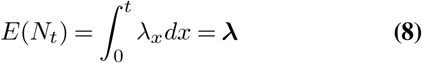

where an estimate 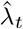 can be made from ***λ*** for all *t* to reconstruct the unobserved process *N*_*t*_.

A similar expression can be constructed for *E*(*Y*_*t*_) = ***µ*** where observed transmissions occur according to a Poisson process with intensity

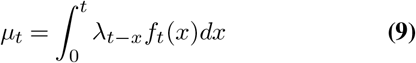

where *f*_*t*_ is the incubation period distribution with infectious period duration *x*. This expression relates the observed number of cases *Y*_*t*_ with the number of infected persons *N*_*t*_, when we assume that the incubation period holds the same distribution regardless of when the individual is infected, and that this intensity arises irrespective of an underlying infection process which independently assigns individuals a specific infectious period^64^.

The previously described convolution is the basis of back projection, where *µ*_*t*_ is assumed to reflect the distribution of symptom onset times, referred to as the epidemic curve, and where the incubation period *f*_*t*_ is assumed to be accurately estimated with sufficient empirical data. To back calculate unobserved infection times, this convolution process is reversed by deconvolving the observed epidemic curve and the incubation period distribution, so that the observed data can reflect past transmission events^63^.

Our approach has some limitations. Eq. (9) does not define a latent period^64^, the time interval between which an individual is infected by a pathogen and when the individual becomes infectious. The latent period often differs from the incubation period, which can only describe the time of infection incidence to the time of symptomatic onset. Therefore the model neglects the impacts of asymptomatic carriers when mapping between infection incidence and case observations. It also fails to take into account transmissions that may occur during an individual’s symptomatic window.

### Incubation Period Distribution

The incubation period distribution (IPD) descibes the delay between individual exposure and symptom onset, viewed as a random unknown variable which is governed by a known probability mass function. Due to the variable nature of SARS-Cov-2 viral interactions with hosts on the population level, the incubation period can vary by many days across individuals, where some hosts may be infectious viral carriers on day 2, or on day 14 after initial infection. Distributions are fitted to data gathered from disease outbreaks, where it is assumed that the data are independently and identically distributed. Following early estimates^65^, we used a generalized gamma distribution with shape and scale parameters 5.807 and 0.948 respectively.

### Convergence of EM method and smoothing step

The EM algorithm described in Results is used for obtaining maximum likelihood estimates in situations where only incomplete data are observed, but a set of ‘complete data’ is defined for which closed-form maximum likelihood estimates exist. Estimates are made by deriving an iterative updating formula, with which we provide an initial estimate for the underlying infection intensities ***λ***, and the Expectation and Maximization steps are simultaneously calculated until convergence.

The EM algorithm converges to one of many configurations maximizing the likelihood, depending on the choice of the 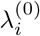 (the values of *λ*_*i*_ used to start the iterations in the EM algorithm). Our initial guess for *λ*_*i*_ is a uniform initialization to avoid biased scaling of the subsequent estimated rates. For each variant, we ensure that the sum of all infection rates is equal to the total number of observations.

After initialization and many iterations which maximize the likelihood of the *λ*_*i*_, a convergence criterion for stopping iterations would typically be based on the value of the likelihood. However, one important difference from a simple EM model is the incorporation of a smoothing step, which will lead to a stopping criterion that is based on the values of the estimated parameters^63^.

After applying our iterating updating equation Eq.(6), let

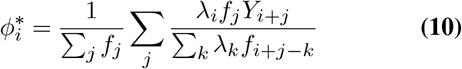

and let

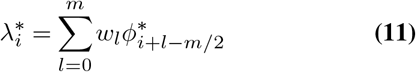

Here, 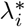 becomes a weighted average of new parameter values produced near *i* during the E and M steps, when applied to old parameter values. The *m* determines the window width for the weighted average. Following previous examples^63,66^ we choose symmetric binomial weights defined as

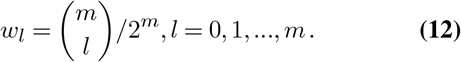

After incorporating this smoothing step into the estimator, it makes sense to choose a convergence criteria based on what we are estimating, since the likelihood is no longer maximized for *λ*_*i*_. Our convergence criterion is then

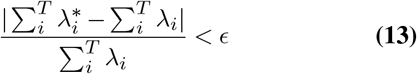

where we used *ϵ* = 10^−4^, and a maximum number of iterations of 250.

### Inference of evolutionary trajectories

Lee et al. describes a method for inferring selection effects using a branching process model developed to represent the transmission of SARS-CoV-2-like viruses, based on observational genomic surveillance data. The estimator for the vector of selection coefficients ***s*** is given by^67^

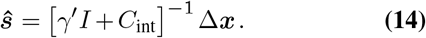

where *γ*^*′*^ is a regularization term proportional to the precision of the prior distribution for the selection coefficients *s*_*i*_, *I* is the identity matrix, *C*_int_ is the covariance matrix of SNV frequencies integrated over time, and Δ***x*** is the change in the SNV frequency vector over time.

### Branching Process

For our simulated branching processes, we chose a constant dispersion parameter *k* = 0.1, with *R* adjusted adaptively such that the number of infected individuals is maintained at around 104 in each “generation.” In real scenarios, *k* and *R* could potentially vary as a function of time, representing different factors (e.g., social distancing) with global effects on disease transmission.

### Regions and Sampling

Regions were selected for back-projected inference if they contained a low to moderate total number of genomes, for computational accessibility. We choose a maximum of 50,000 sequences per region to allow for a reasonable computation time for the estimation of back-projected infection weights, and for further processing steps such as the calculation of the region’s covariance matrix. As in prior work, we also required a minimum of at least 200 sequences and 15 days of data for a region to be included, reasoning that extremely short and/or poorly sampled trajectories would not be informative.

We prioritized analyzing regions where sampling distributions displayed data gaps over time regions, especially between outbreaks of variants (i.e., regions that were poorly sampled between selective sweeps of VOCs), over which back-projection could act to smooth over. We also prioritized regions with large fluctuations in sampling, where we anticipate that smoothing from back-projection will be especially useful.

### Processing Pipeline

The steps taken for data processing are similar to those written in Lee et al., with some modifications.

Pre-processing steps:

1. We removed incomplete sequences with excess gaps or ambiguous nucleotides in more than 1% of the genome that do not correspond to known deletions. We also remove sites where gaps appeared in over 90% of cases, as these likely represent rare insertions or sequencing/alignment errors.
2. We separated sequence data into regions and then ordered time series in those regions.
3. We performed imputation for sites with ambiguous nucleotides or unexpected gaps with a maximum gap frequency threshold of 50%, to separate likely deletion events from poor quality data.

Back-projection is implemented after sequences are preprocessed. The steps required for the method are:

1. Separating sequences into the 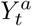. We divide our data into sets of “non-unique” sequences that appear more than once in a region. Sequences that appear one time will have the same corresponding 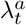 after performing the deconvolution, determined by the incubation period distribution.
2. Back-projection is performed across 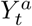, and 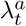 are generated. 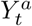 gives estimated infection *weights*, compared to observed sequence *counts* in *Y* ^*a*^.

After performing back-projection, we eliminated time periods with poor sampling by dropping times where the sum of all sequence weights within a sliding 5 day window was less than 20. These criteria are analogous to those described by Lee et al., which required a minimum of 20 sequences per 5 day period. To compare with results from the Lee et al. approach, we restricted our analyses to SNVs observed at frequencies above 5% without backprojection (see **Supplementary Fig. 19**).

After the back-projected dataset and the non-back-projected comparison set are processed, we then can compute SNV frequency changes and their covariance matrices for each region and integrate them over time. Selection coefficients are inferred by inverting the integrated covariance matrix. In these computation steps, counts as integers are replaced by weights as floats in the back-projected case.

### Data and code

Processed data and code used in our analysis are available in the GitHub repository located at https://github.com/bartonlab/paper-epi-backprojection. This repository also contains Jupyter notebooks that can be run to reproduce the results presented here.

---

**Supplementary Fig. 1.**
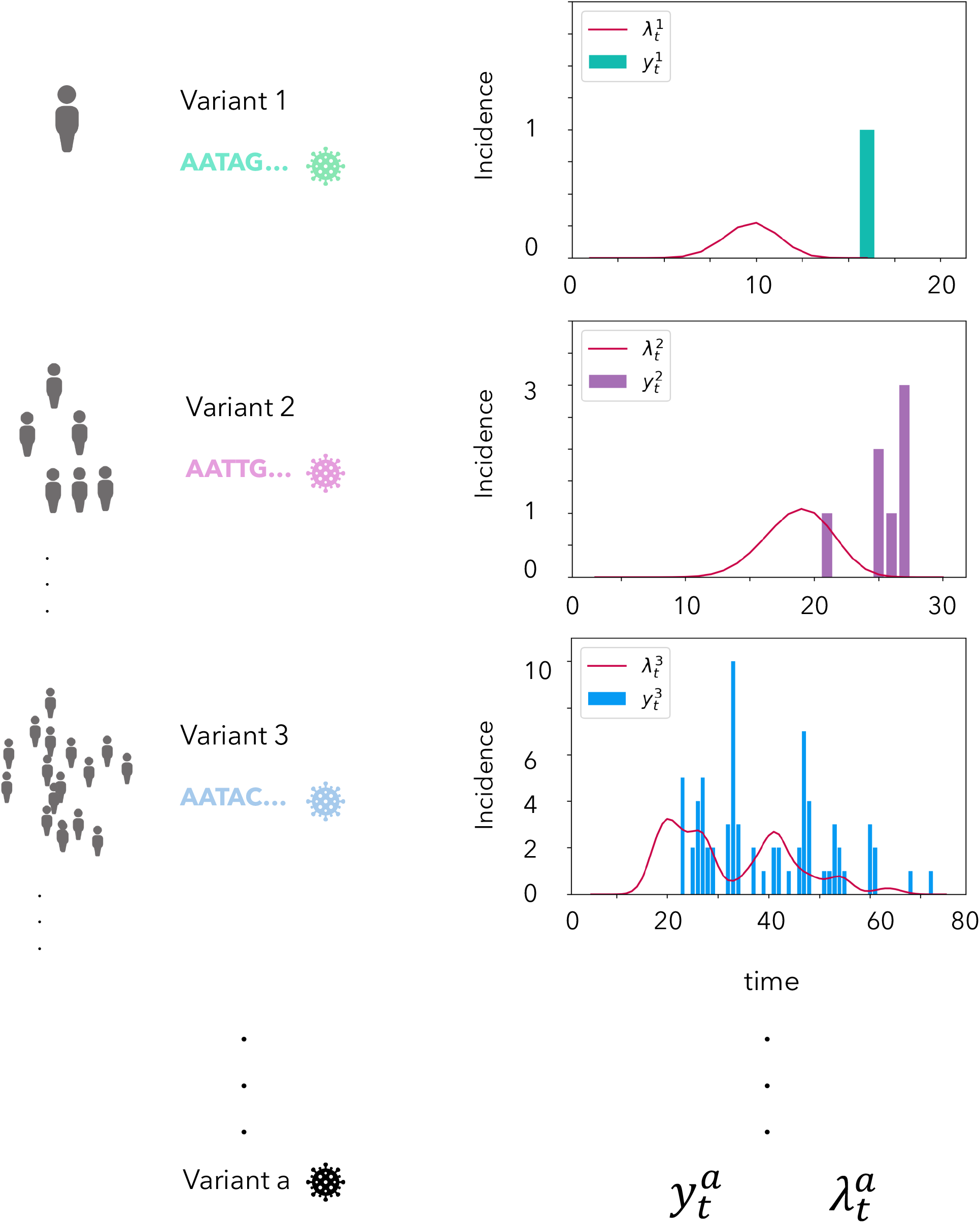
Dividing up sequences for back-projection processing, graphical illustration. The SARS-CoV-2 genome is of size 30kbp (kilo base pairs). For a viral genome this large, some SNV (single nucleotide variant) mutations only appear in combination once in a regional outbreak. Other SNVs in the same combination spread to multiple people during an outbreak. We group individuals passing along the same genetic variation of SARS-CoV-2 before smoothing these sequence counts via back-projection. We perform this division as a pre-processing step to allow us to replace integer value sequence counts in our additive model of evolution and selection inference, with normalized non-integer weights of counts at a given discrete time point. Each SNV, *a*, is separated into its own population of observed cases, 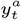, with which we estimate the incidence 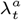 (Methods). In both our original method and our implementation of back-projection, these “populations” of unique and non-unique sequences have their alleles combined with an additive model, in order to create single allele frequencies which make up that allele’s evolutionary history.

**Supplementary Fig. 2.**
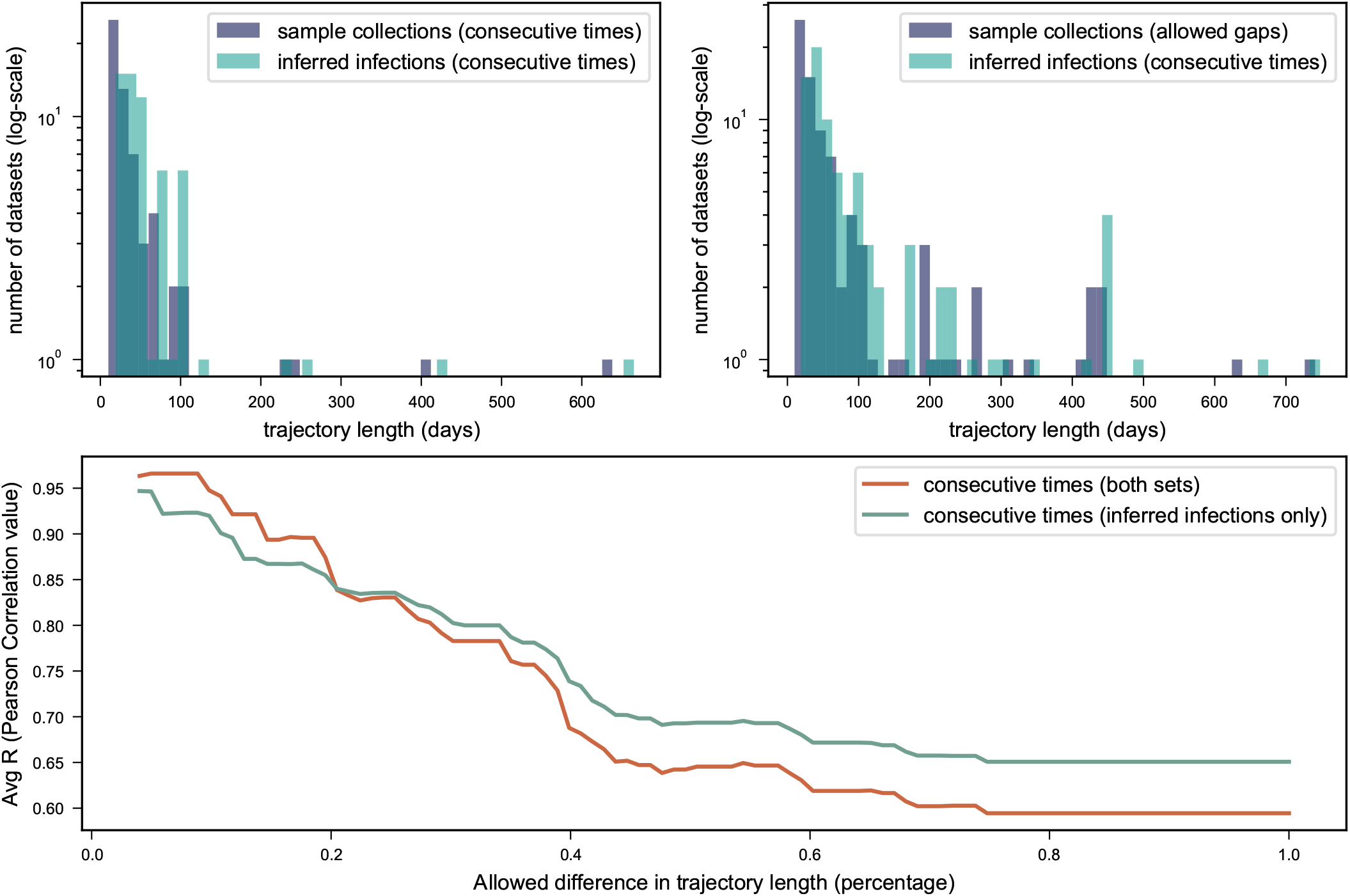
Pearson’s correlations at different trajectory length thresholds. As a result of back projection increasing trajectory length and providing smoothed estimates over data gaps, the correlations we obtain when we compare the inferred selection effects of back projected inferred infections, and unprocessed sample collections, can vary. In particular, the difference in lengths between the starting trajectory and the back-projected trajectory can have an effect on the Pearson correlation values that we calculate. Figure **a**. and figure **b**. show log-scaled histograms of the length of trajectories, in both a set of completely contiguous trajectories for both data, and a set where back-projected estimates have contiguous data, but we allow for the sample collections to contain data gaps. For both sets, we calculate the average Pearson Correlation based on a threshold which subsets the allowed trajectories to be averaged, based on how dissimilar the length of the inferred infection trajectories are from the sample collection trajectories. We obtain good correlation (0.97-0.85) for paired trajectory lengths that do not deviate by more than 20 %. From there, as the length of the inferred infection trajectories becomes more dissimilar to the sample collection trajectories, the average correlation falls off.

**Supplementary Fig. 3.**
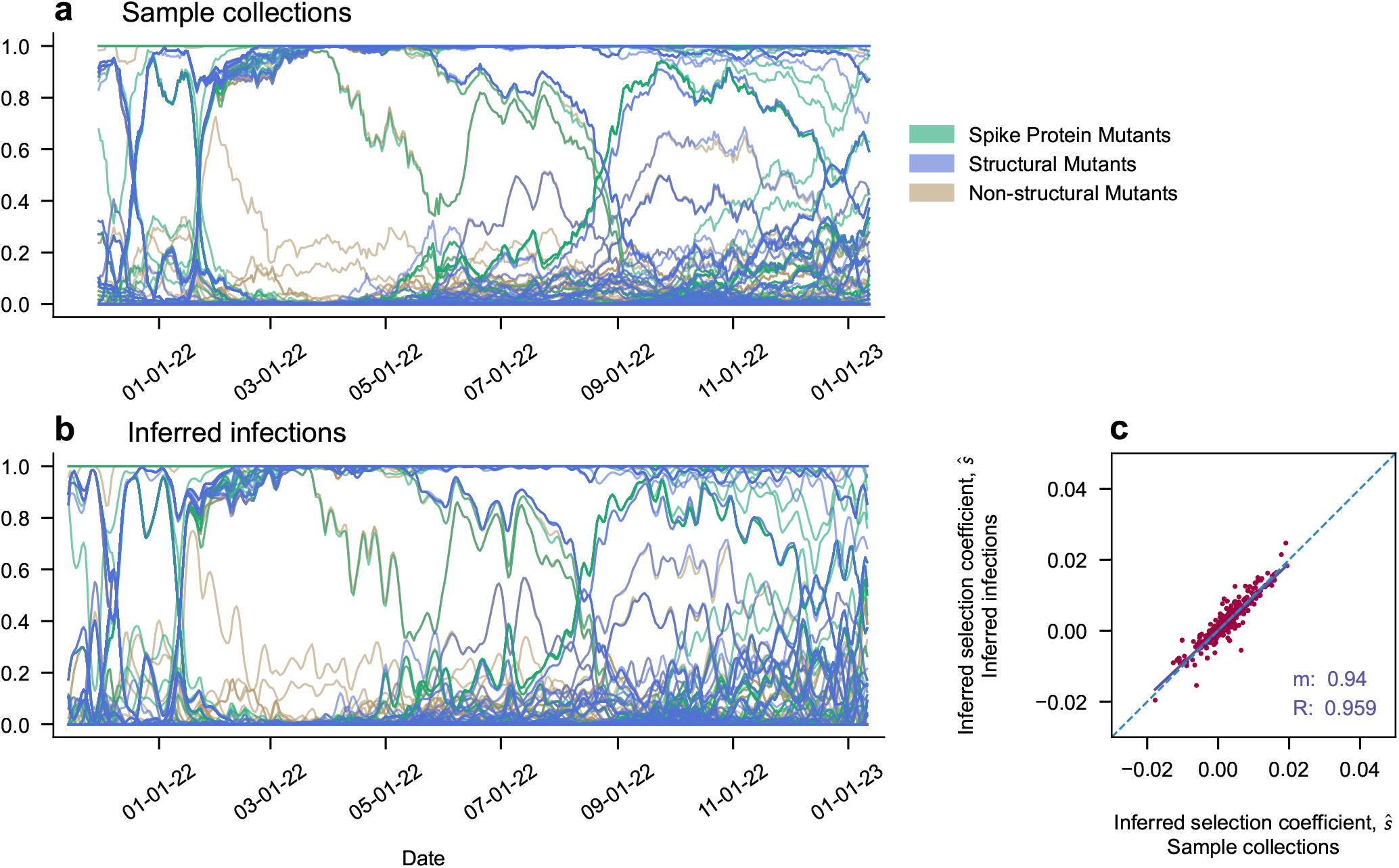
Selection coefficients with and without back-projection are highly correlated in long trajectories (Hong Kong). Trajectories across long time periods show smoother dynamics, with large agreement.

**Supplementary Fig. 4.**
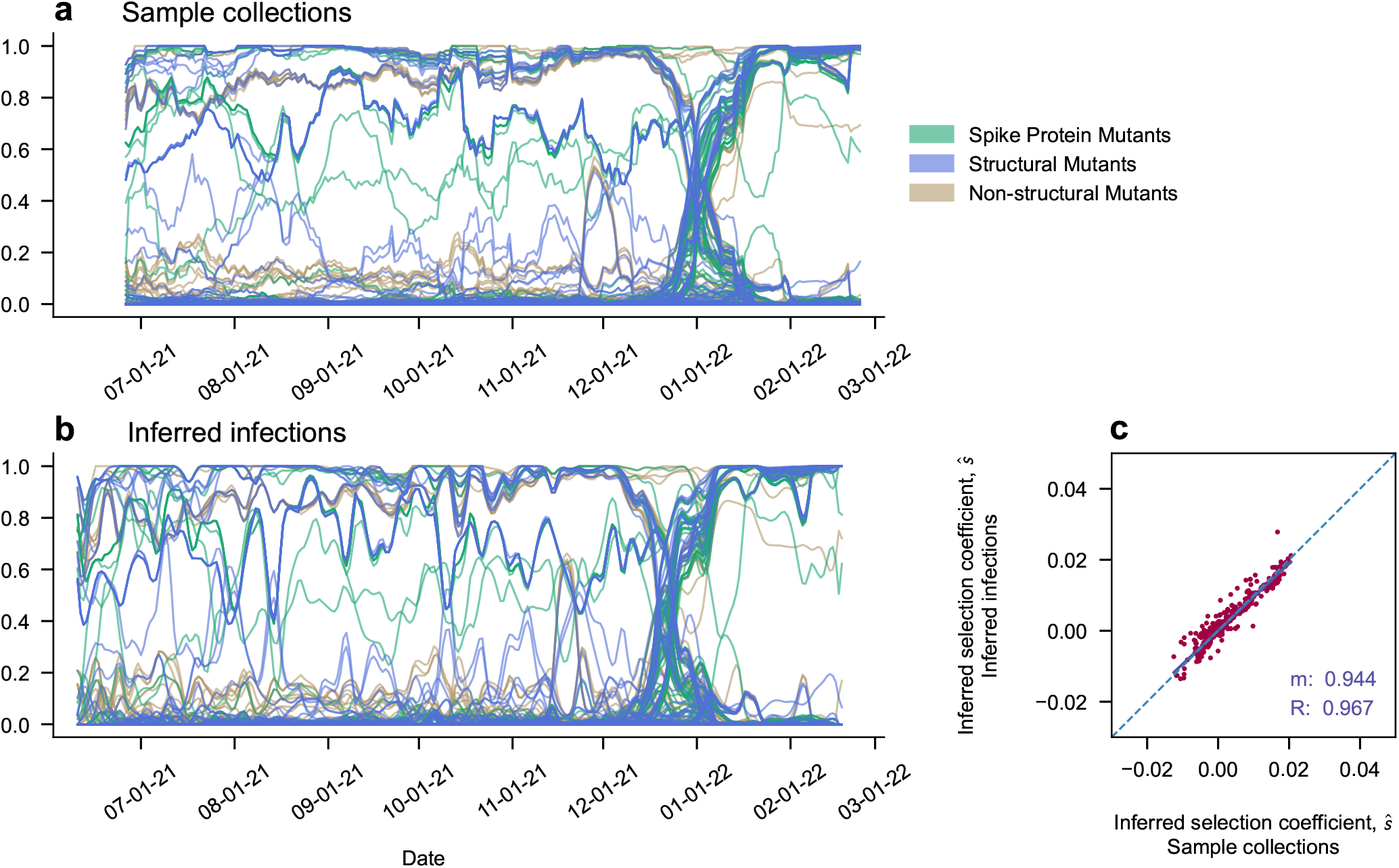
Selection coefficients with and without back-projection are highly correlated in long trajectories (Karnataka). Trajectories across long time periods show smoother dynamics, with large agreement.

**Supplementary Fig. 5.**
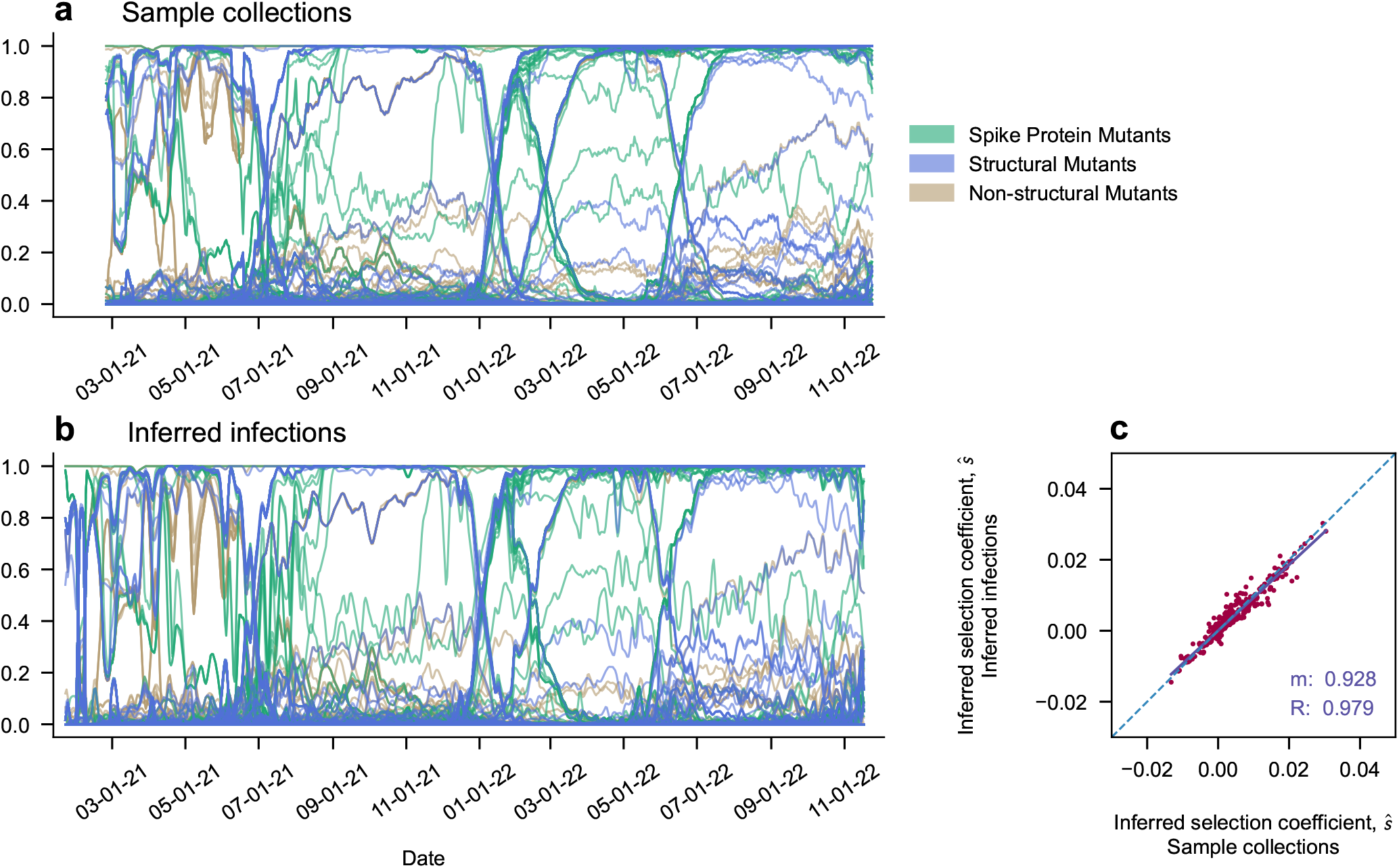
Selection coefficients with and without back-projection are highly correlated in long trajectories (Slovakia). Trajectories across long time periods show smoother dynamics, with large agreement.

**Supplementary Fig. 6.**
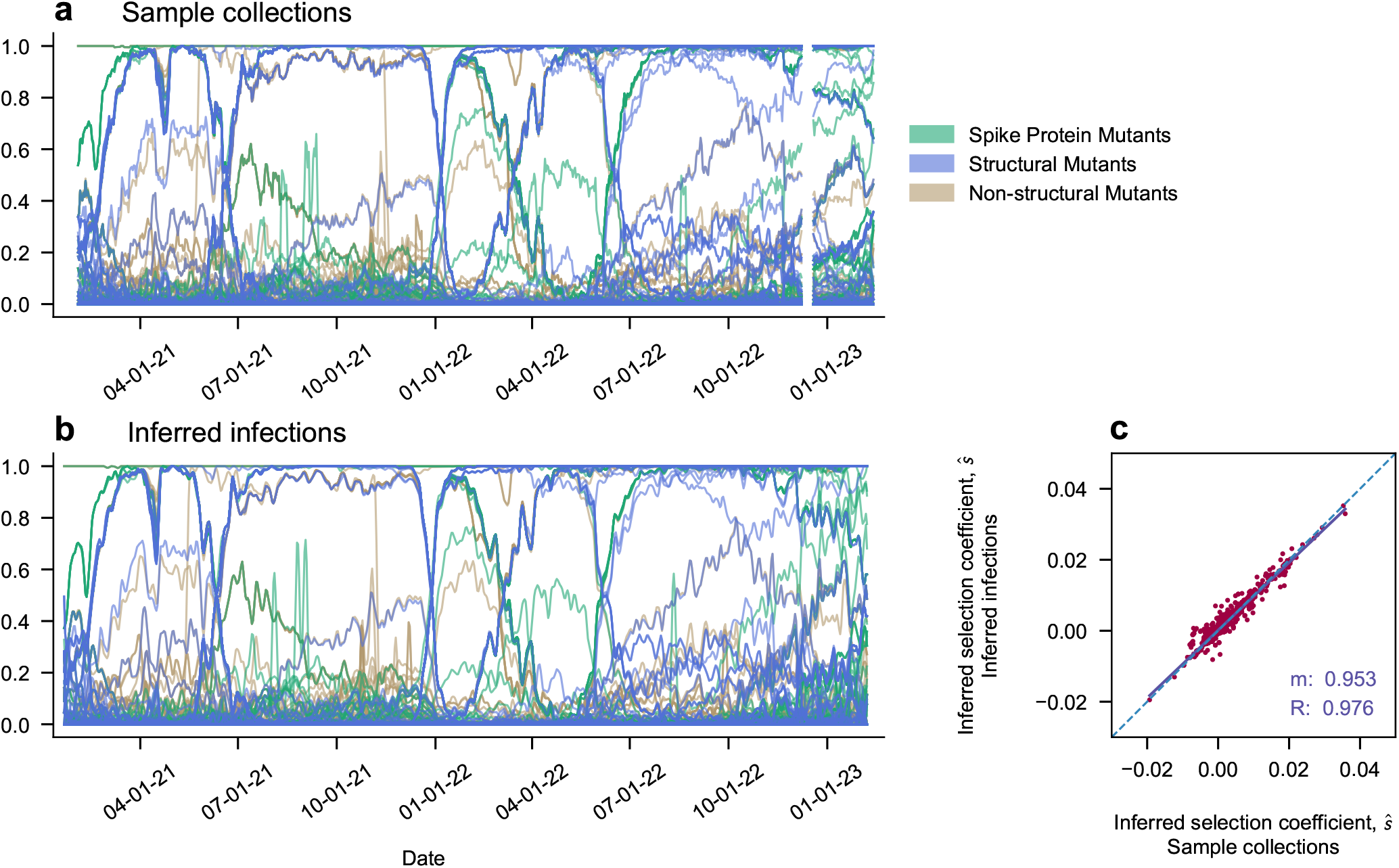
Selection coefficients with and without back-projection are highly correlated in long trajectories (Croatia). Trajectories across long time periods show smoother dynamics, with agreement even allowing for small gaps in the unprocessed frame.

**Supplementary Fig. 7.**
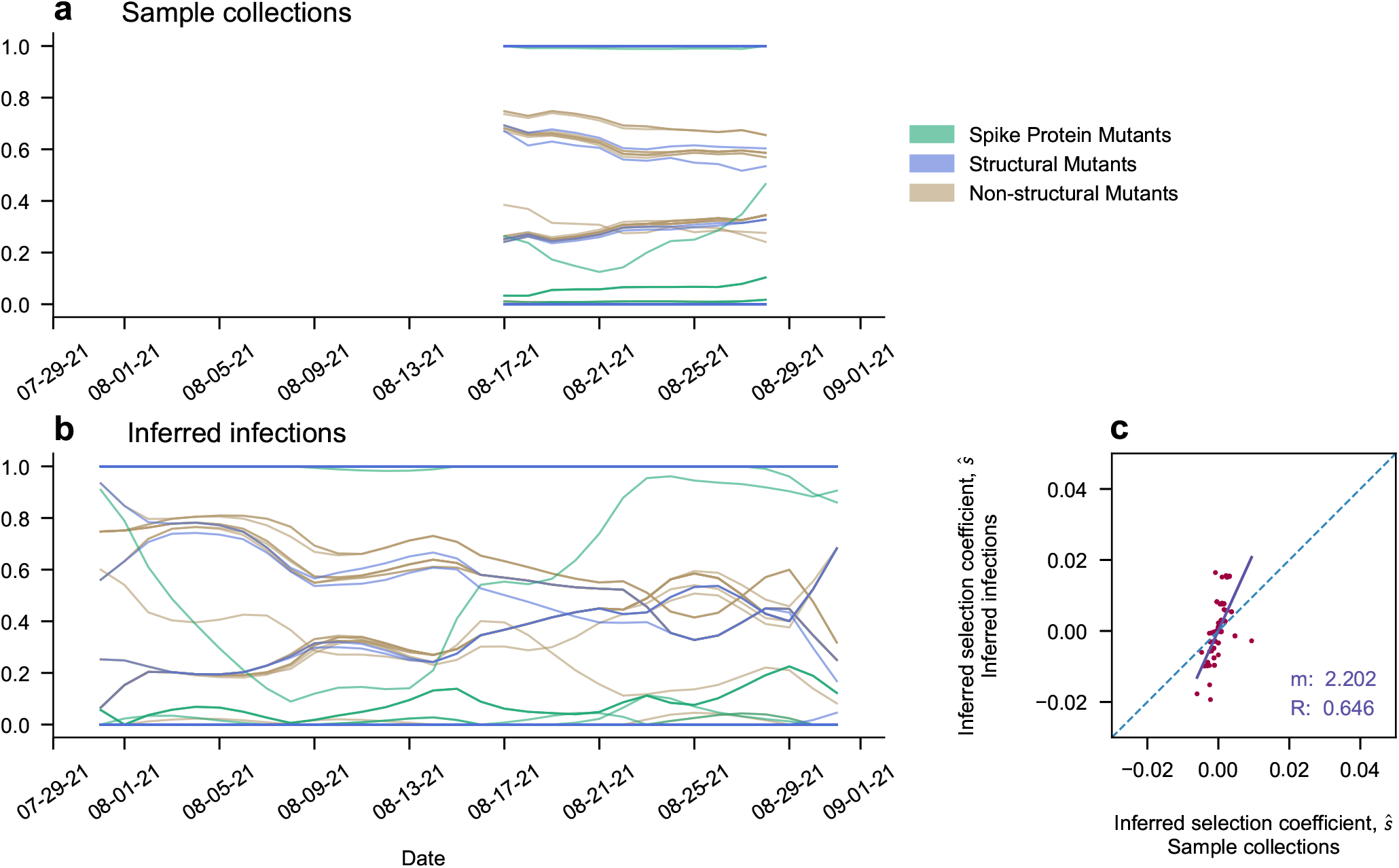
Correlated effects can vary when comparing trajectories of different lengths (Kenya). Back-projected estimates often provide a greater amount of trajectory information

**Supplementary Fig. 8.**
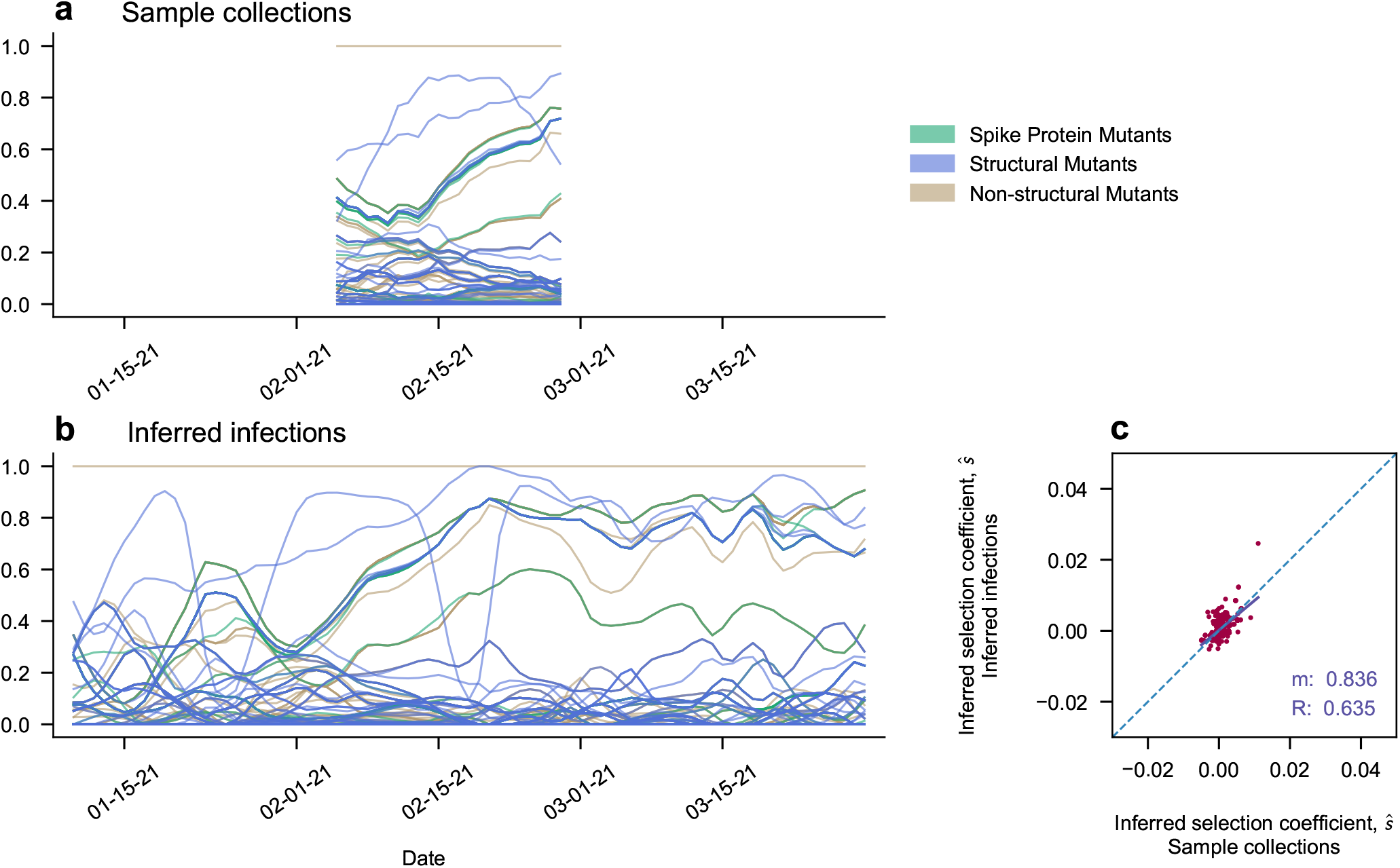
Correlated effects can vary when comparing trajectories of different lengths (Reunion). Back-projected estimates often provide a greater amount of trajectory information

**Supplementary Fig. 9.**
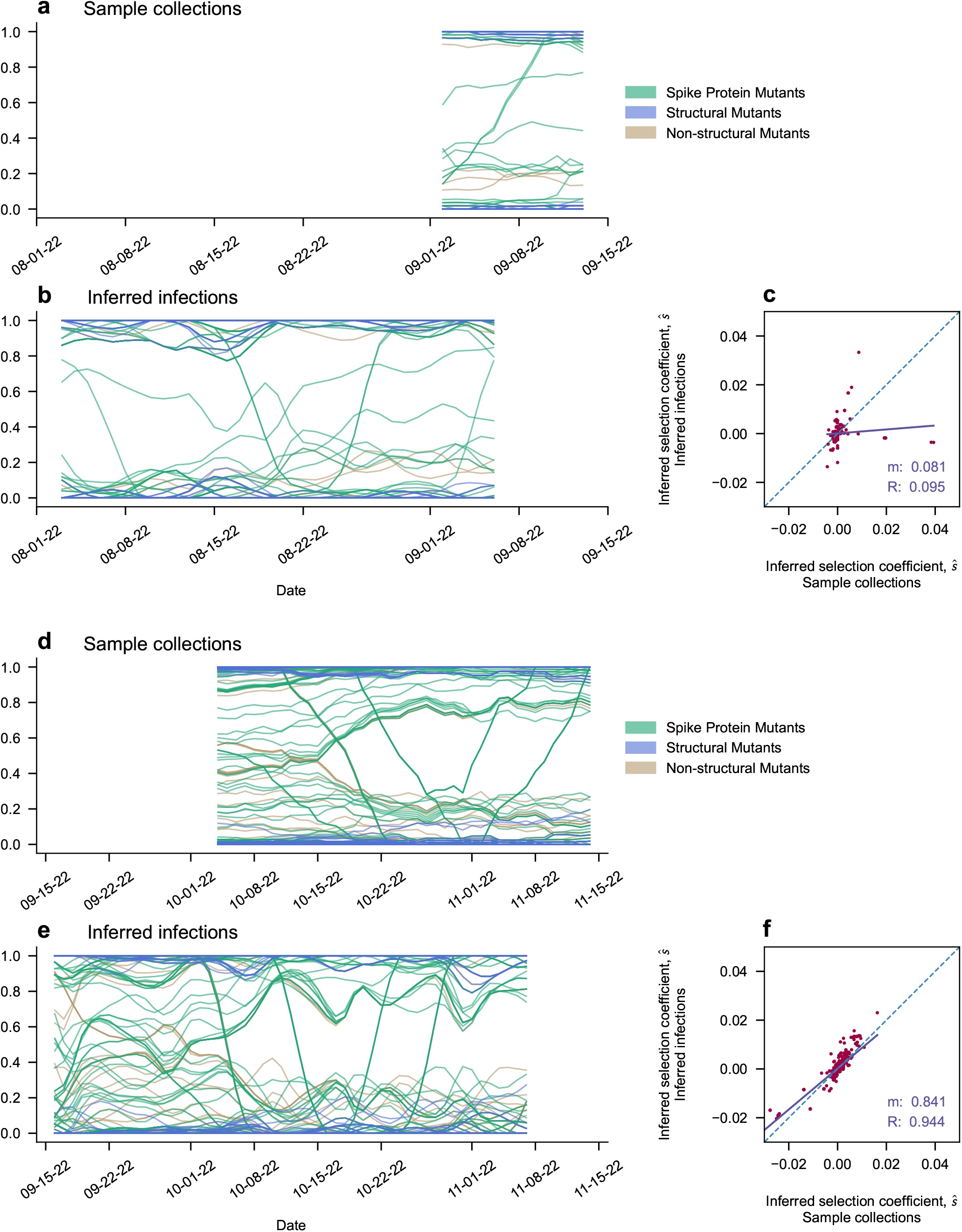
Correlated effects can vary when comparing trajectories of different lengths (Gujarat). Back Projected estimates often provide a greater amount of trajectory information. Here, the displacement of Omicron BA.1 by more transmissible variants such as BA.2, BM.1, BA.5, BQ.1 and XBB.1 cause sweeps of Spike protein mutations.

**Supplementary Fig. 10.**
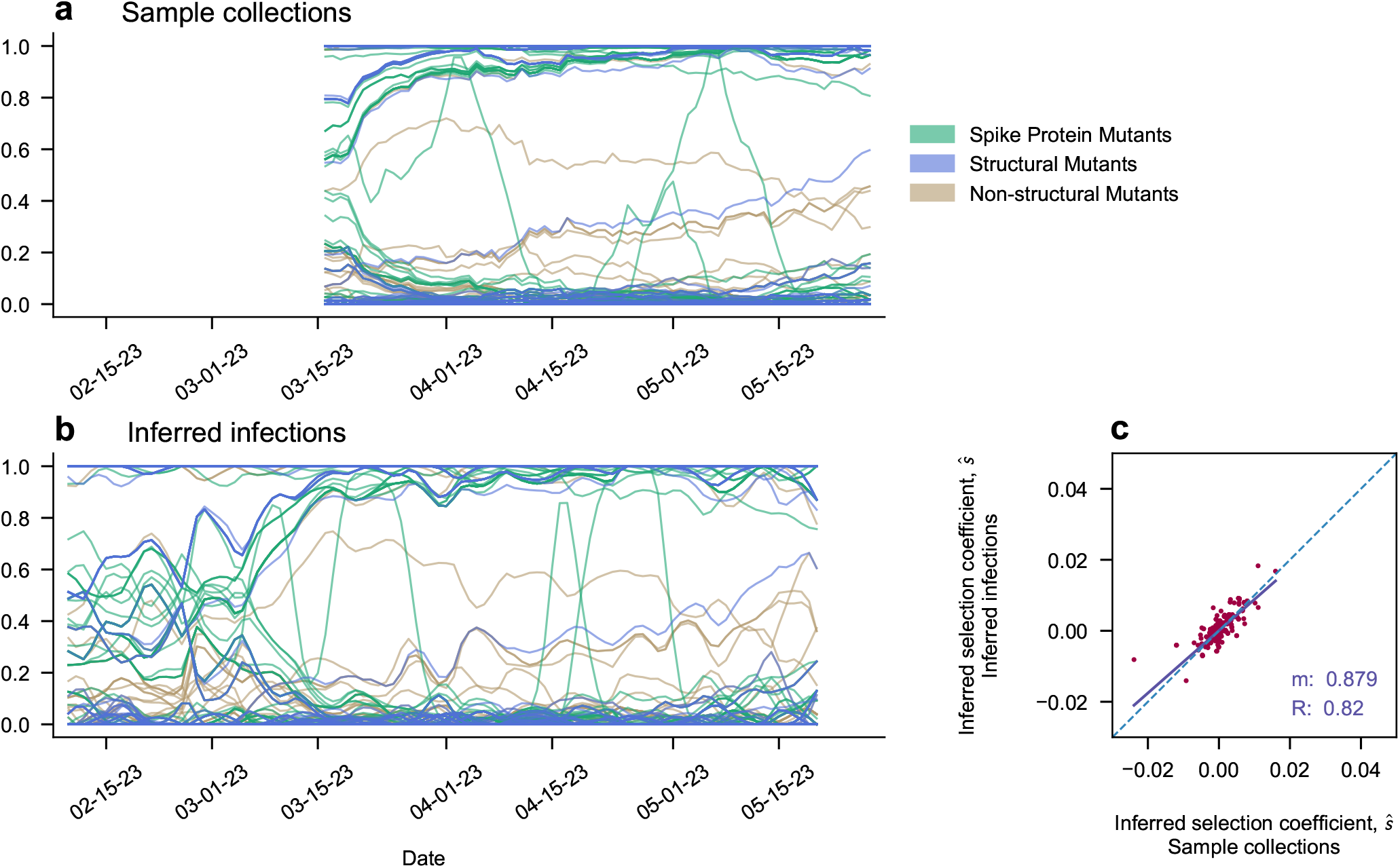
Correlated effects can vary when comparing trajectories of different lengths (Croatia). Back Projected estimates often provide a greater amount of trajectory information. Here, the displacement of Omicron BA.1 by more transmissible variants such as BA.5, BQ.1 and XBB.1 cause sweeps of Spike protein mutations.^68^

**Supplementary Fig. 11.**
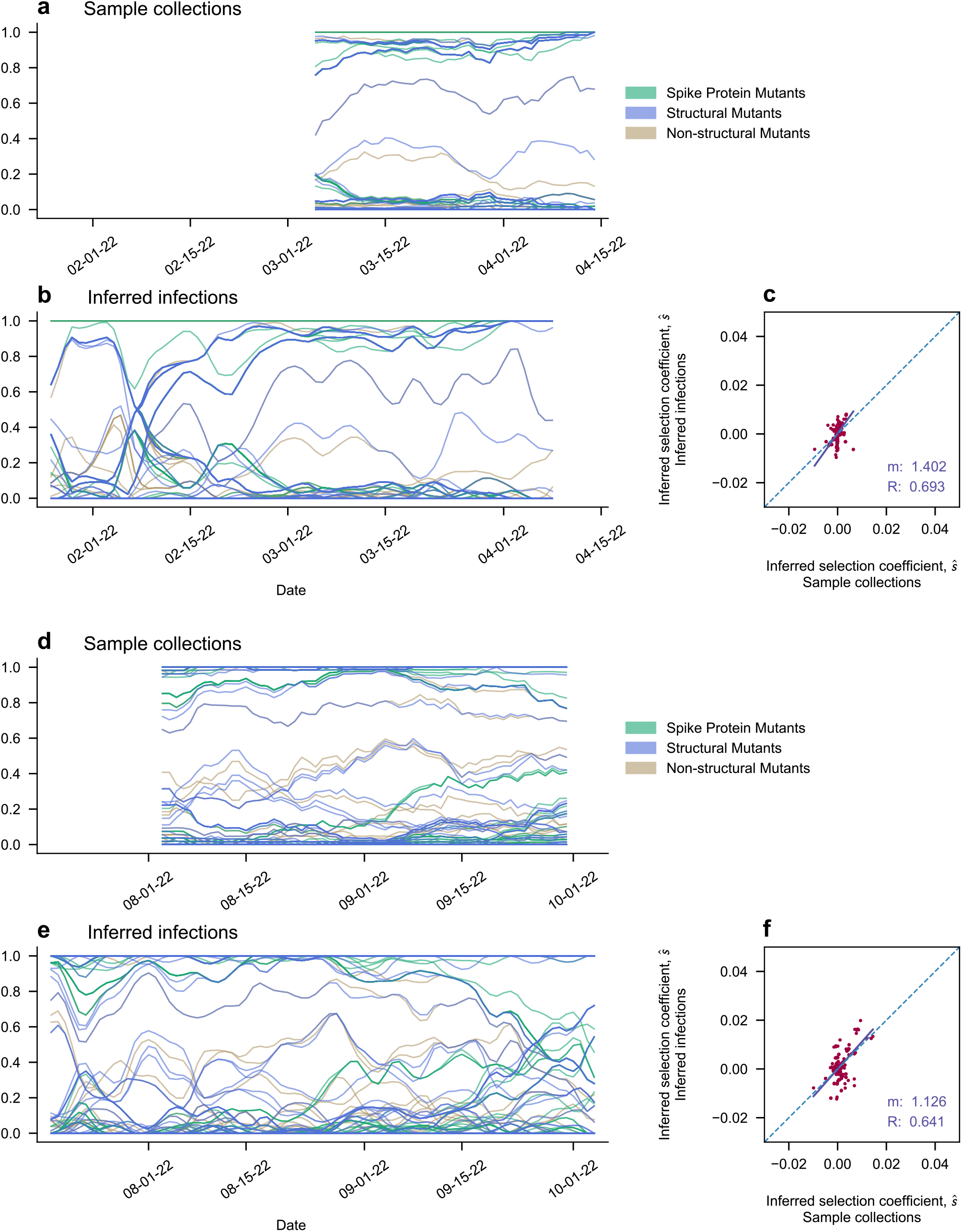
Outbreak: Vietnam. Trajectory dynamics throughout waves of Omicron in Vietnam. Omicron appeared in Vietnam in early 2022^69^ and by late 2022 had many circulating variants of Omicron subtypes.^70^

**Supplementary Fig. 12.**
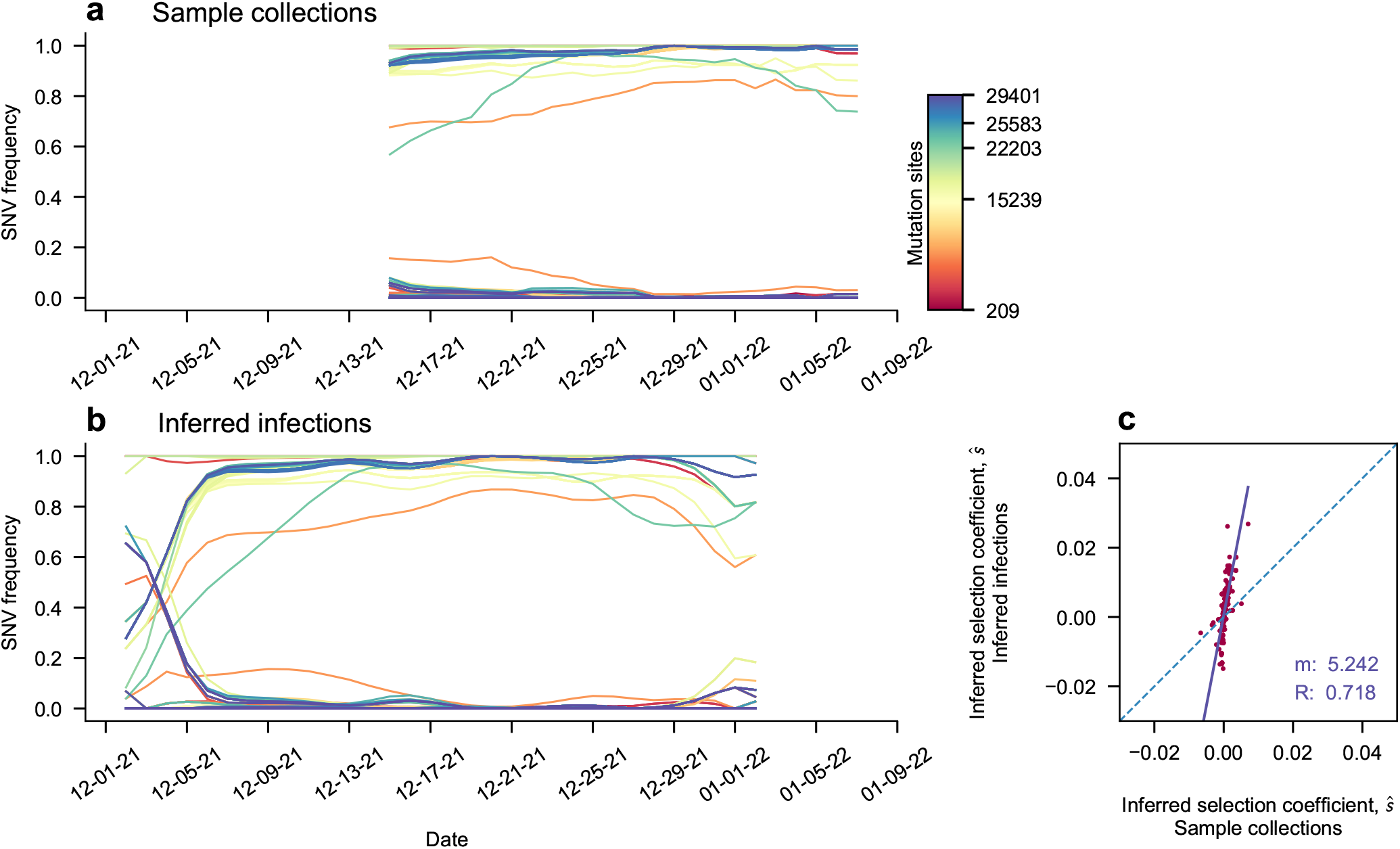
Missed rise of Omicron sweep in multiple regions (Kenya). Some regions contain poor sampling before the Omicron BA.1 rise. Back-projecting trajectory information provides additional information to infer stronger selection coefficients.

**Supplementary Fig. 13.**
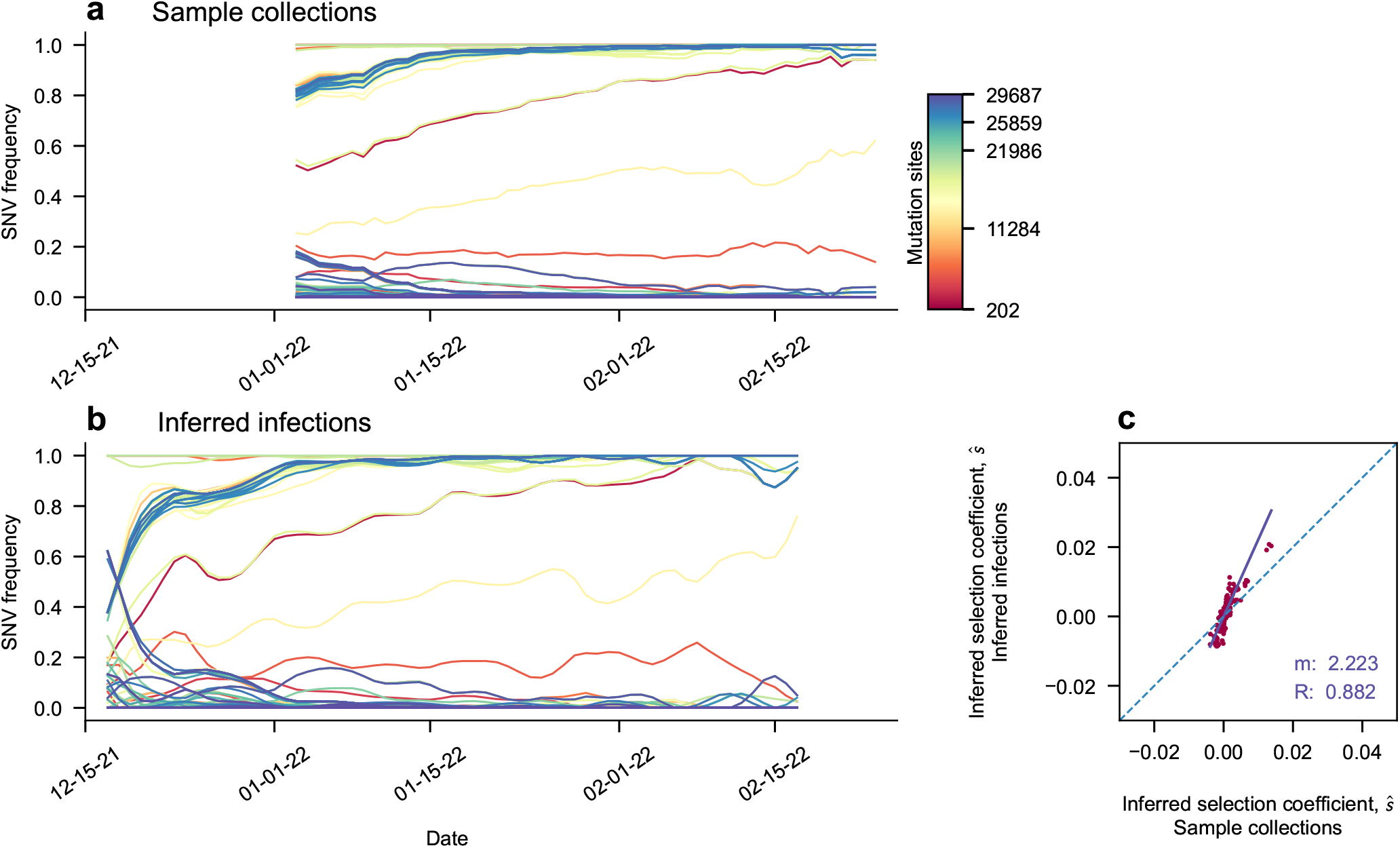
Missed rise of Omicron sweep in multiple regions (Wyoming). Some regions contain poor sampling before the Omicron BA.1 rise. Back-projecting trajectory information provides additional information to infer stronger selection coefficients.

**Supplementary Fig. 14.**
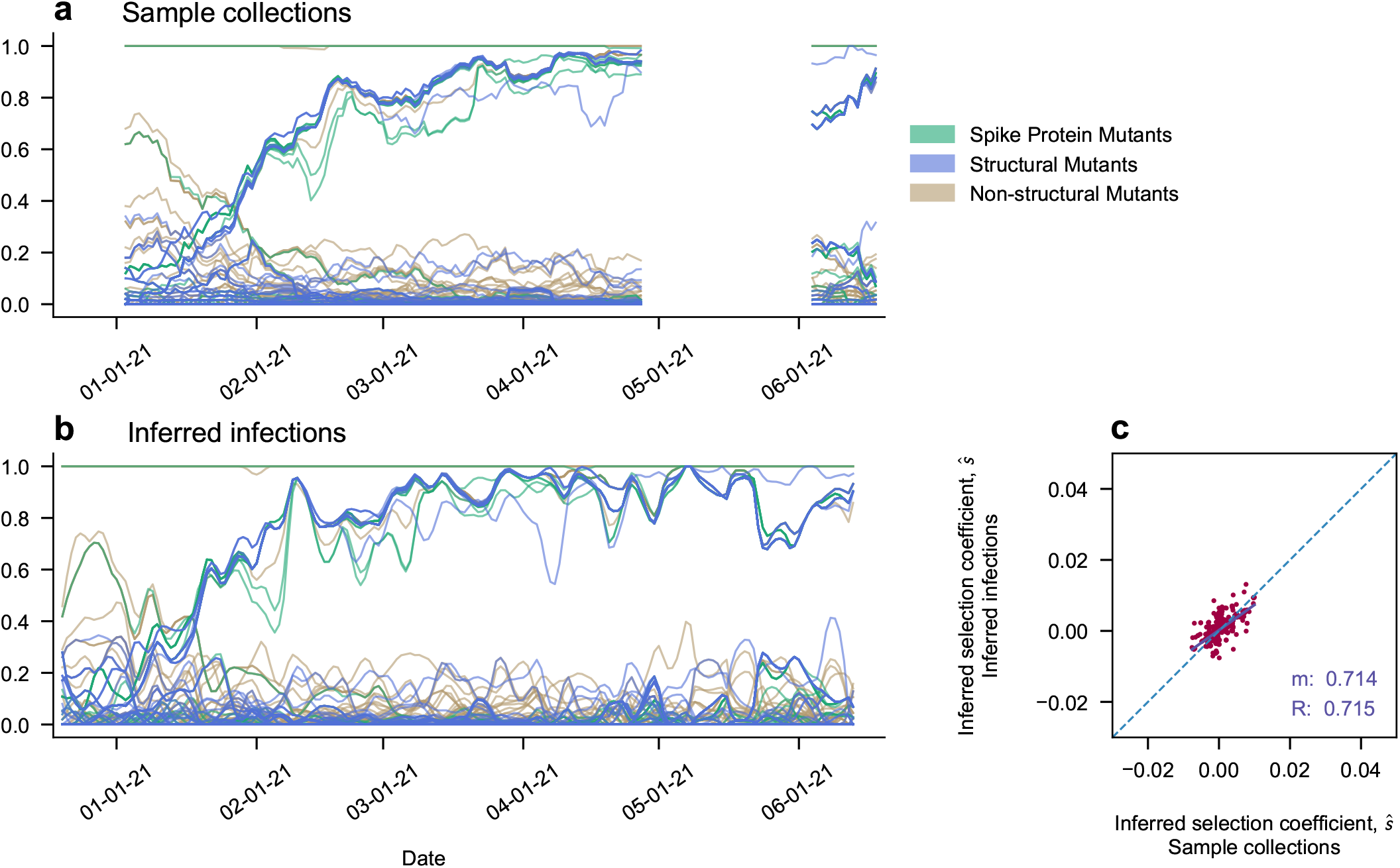
Back-projected estimates smooth over gaps in data (Romania). Back-projected trajectories fill in gaps in data.

**Supplementary Fig. 15.**
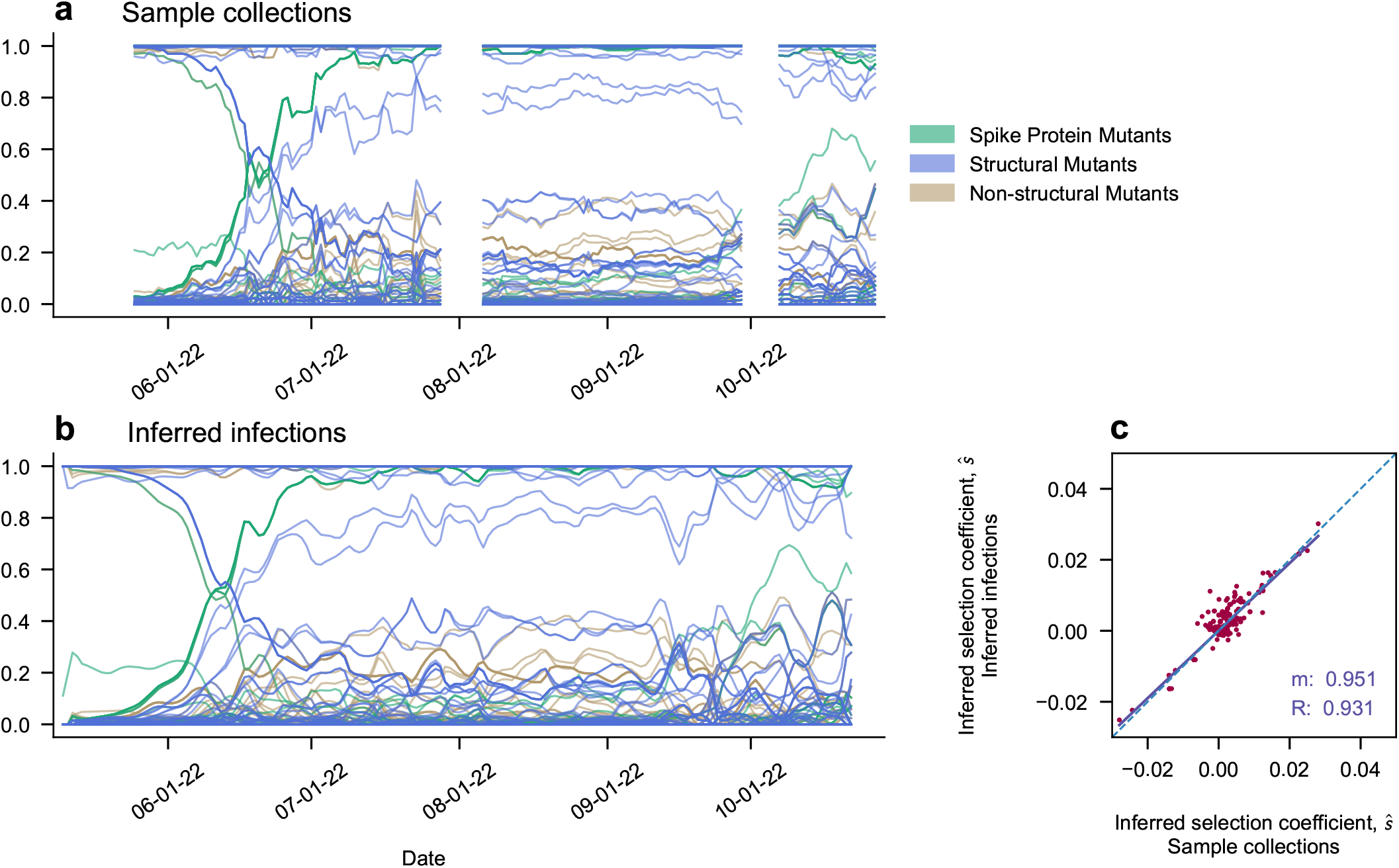
Back-projected estimates smooth over gaps in data (Reunion). Inferred trajectories fill in gaps in data.

**Supplementary Fig. 16.**
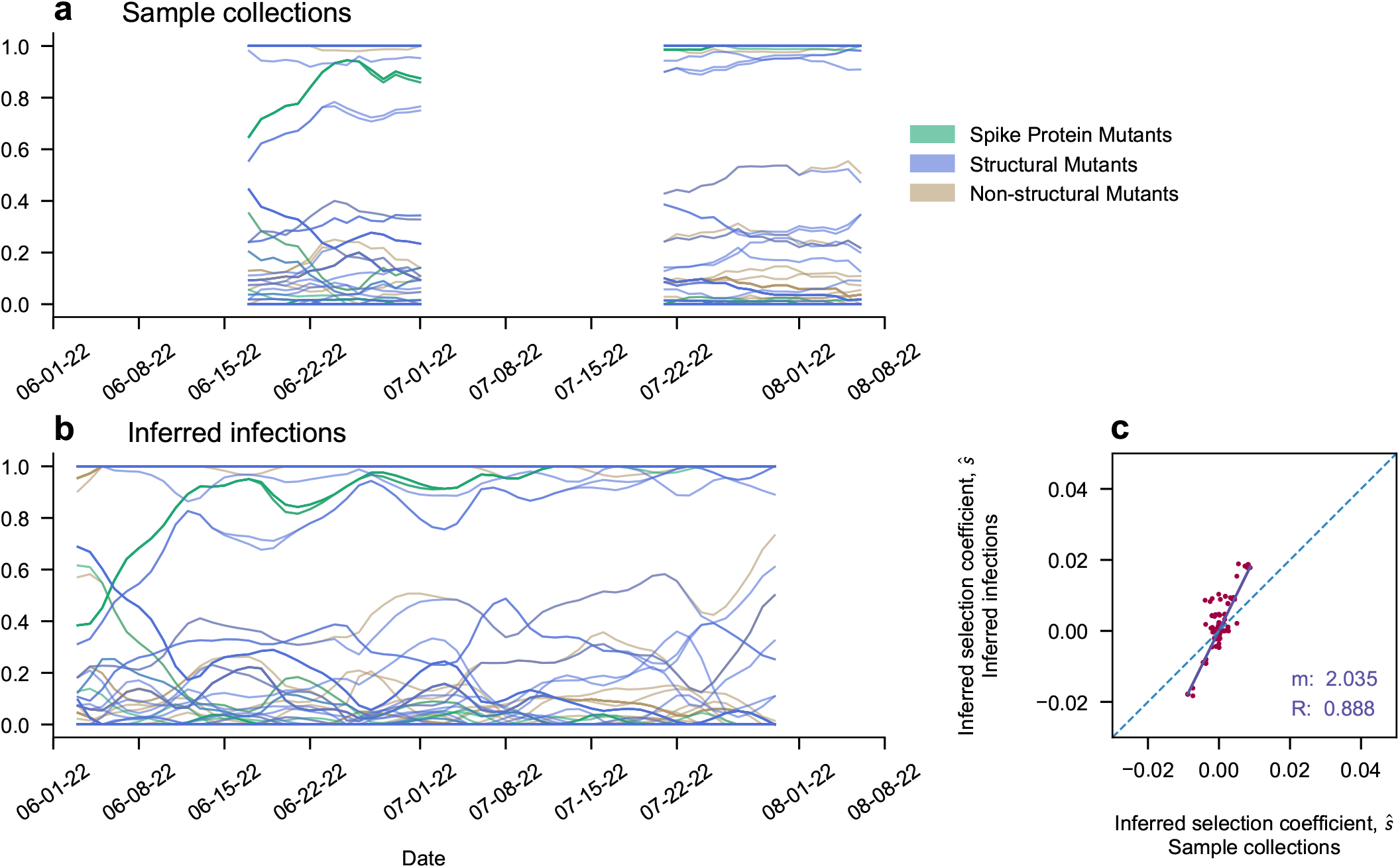
Back-projected estimates smooth over gaps in data (Cyprus). Inferred trajectories fill in gaps in data.

**Supplementary Fig. 17.**
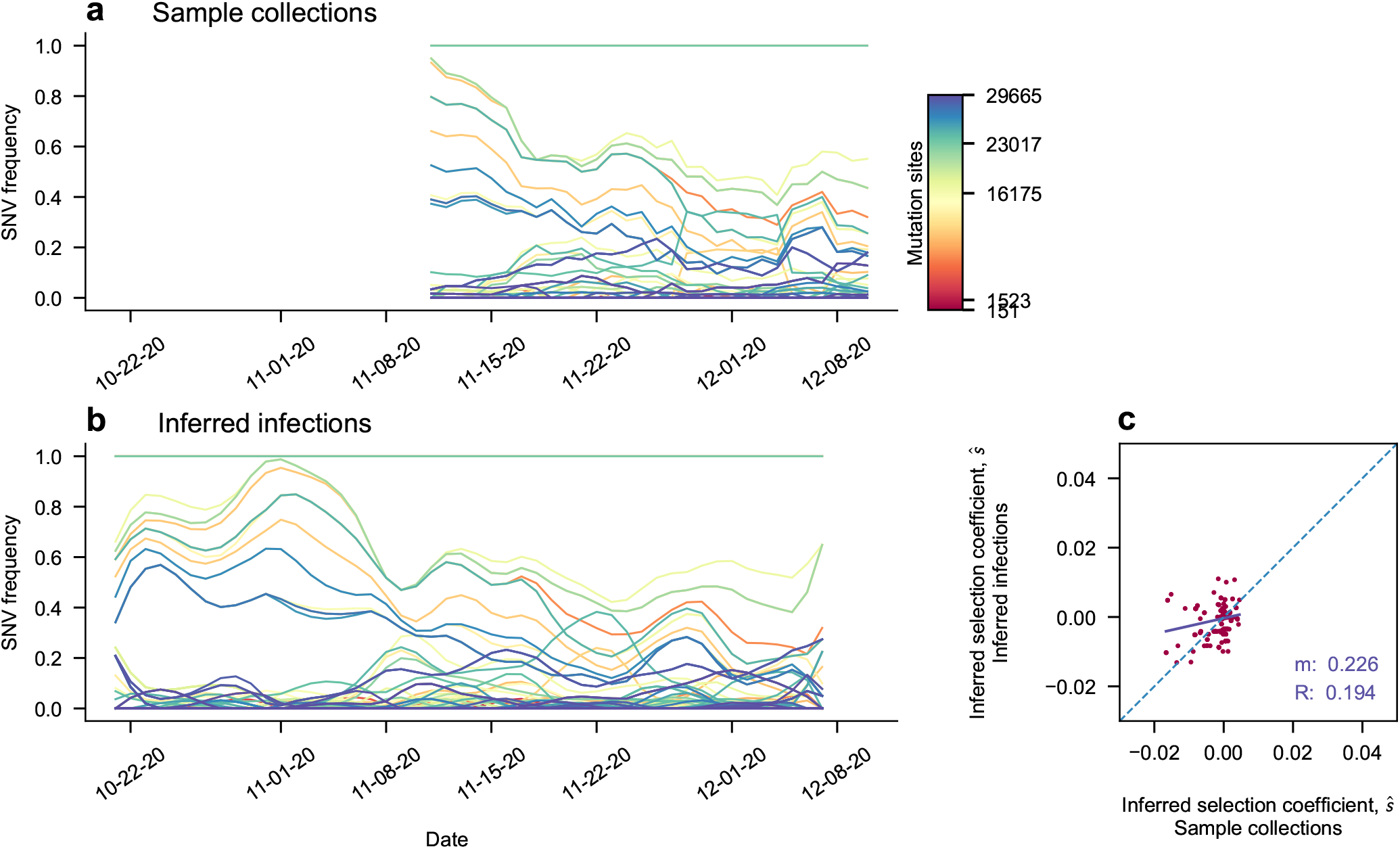
Increased sampling at the beginning and end of an outbreak (Romania). Significant increases in trajectory length with back-projection lead to more moderate correlations between selection coefficients inferred with and without back-projection due to differences in information about mutations driving changes in transmission.

**Supplementary Fig. 18.**
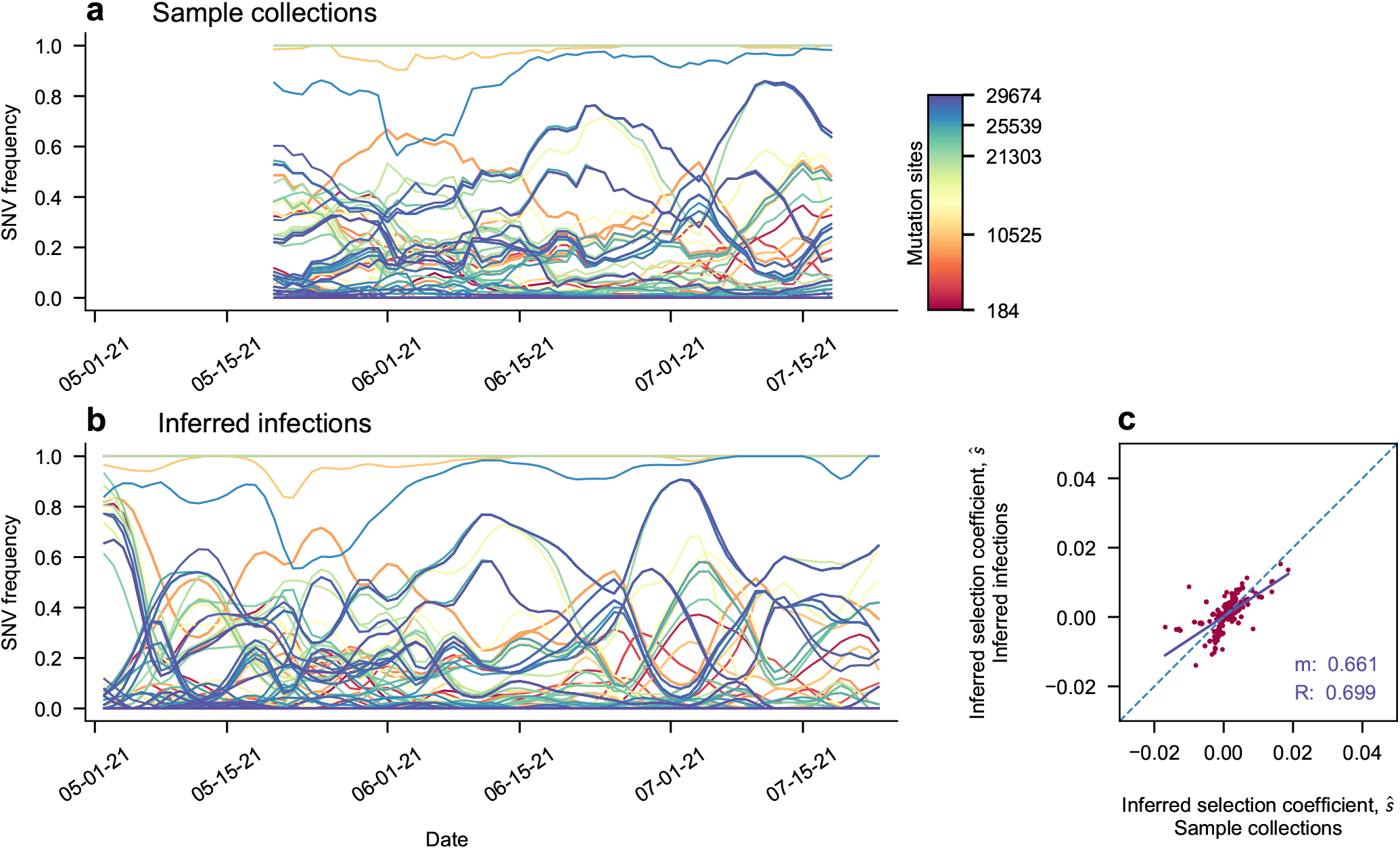
Increased sampling at the beginning and end of an outbreak (Manitoba). Significant increases in trajectory length with back-projection lead to more moderate correlations between selection coefficients inferred with and without back-projection due to differences in information about mutations driving changes in transmission.

**Supplementary Fig. 19.**
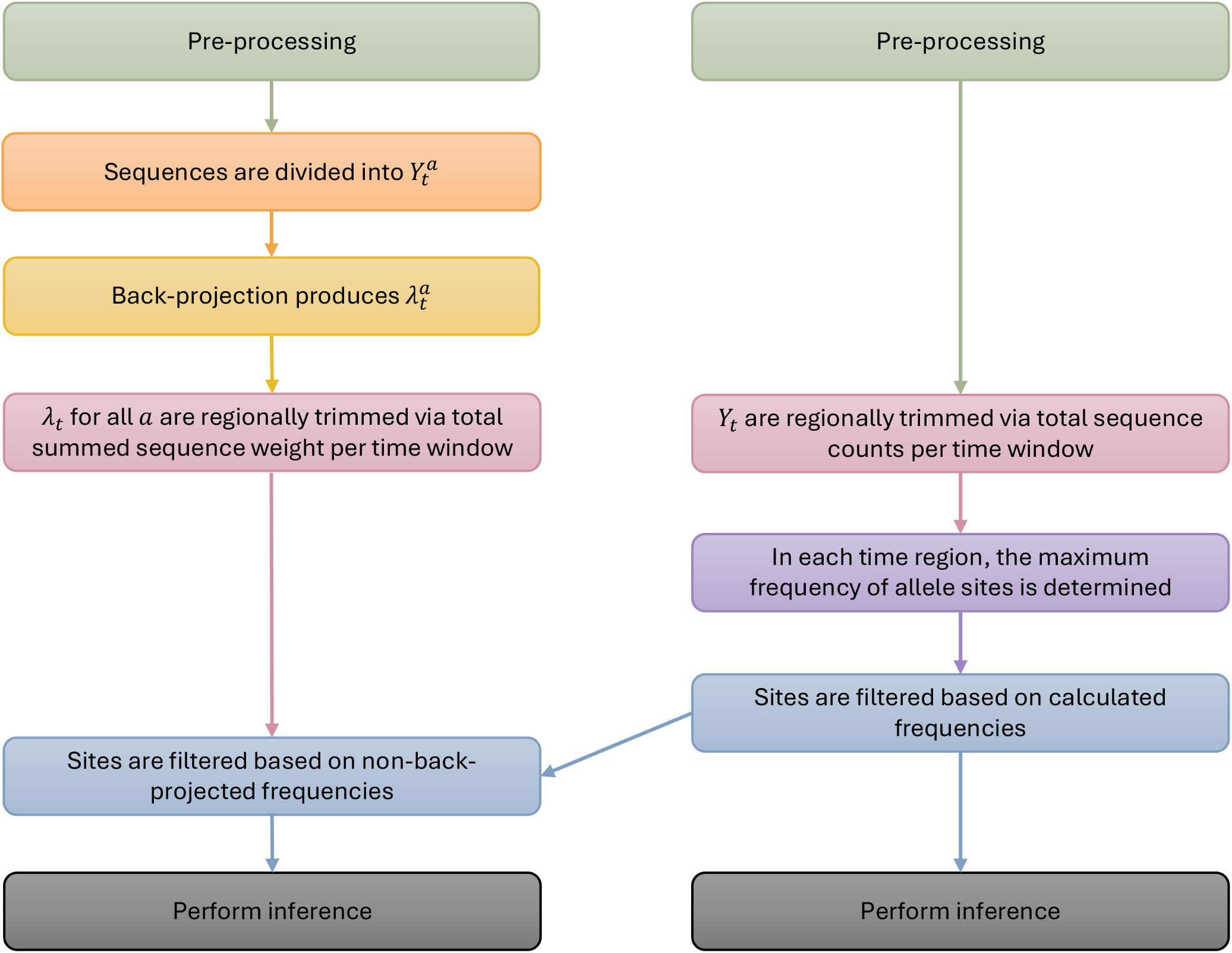
Processing steps with and without back-projection. For comparison, we perform analysis with and without the back projection steps. To allow comparison, some processing steps are performed with the same criteria, as the trimming step, and others are adapted to depend on the parameters in the unprocessed set. In particular, a filtering step chosen to eliminate rare mutations is performed in the unprocessed set, and those same filtered alleles are chosen for comparison in the back-projected dataset. Code for filtering back projected alleles is available, but not used in this analysis.

